# Flow Virometry-Based Profiling of Viral Population Diversity

**DOI:** 10.64898/2026.11.18.767602

**Authors:** Tongyu Liu, Nick Berg, Callie J. Donahue, Steven D. Planitzer, Antonio Santos-Peral, Edward A. Partlow, Balindile B. Motsa, Tijana Ivanovic

**Affiliations:** Laboratory of Viral Diseases, National Institute of Allergy and Infectious Diseases, National Institutes of Health, Bethesda, MD 20892, USA; Department of Biochemistry and Biophysics, Brandeis University, Waltham, MA 02453, USA

**Author notes:** These authors contributed equally to this work (Tongyu Liu, Nick Berg, Callie J. Donahue, Steven D. Planitzer).

## Abstract

Viruses exhibit great phenotypic heterogeneity. Flow virometry enables high-throughput analysis of phenotypic diversity in viral populations, while flow-based sorting allows subpopulations to be isolated for separate characterization. Despite their advantages, flow virometry and sorting have not yet been widely adopted in virology partly because of limited protocols and underexplored applications. Here, we provide a practical guide for analyzing and quantifying virus particles from diverse families using flow virometry enhanced by fluorescence labeling. We further explore applications beyond basic and static characterization of virus populations. We demonstrate that flow virometry can report on interactions between virions and binding partners, as well as between virions. It provides a particle denominator that, combined with genome copy measurements, enables estimation of viral population ploidy. We also show how dynamic processes, such as viral membrane fusion, can be measured in real time. Finally, we demonstrate the feasibility of sorting virions based on their intrinsic properties, such as morphology, for downstream study. This practical guide provides a roadmap for wider implementation of flow virometry in virology, while the applications illustrated here demonstrate potential to address a variety of questions about virion properties, interactions, and dynamics at the single-particle level.

## Introduction

Virus populations are diverse at both genetic and phenotypic levels. While genetic diversity has been extensively studied through advances in sequencing, individual virions also vary in genome packaging and organization, particle morphology, and maturation state, with important biological consequences. For influenza A virus (IAV), particle morphology affects susceptibility to antibody neutralization, with filamentous morphs remaining infectious under conditions that neutralize spherical virions (1, 2). For retroviruses like human immunodeficiency virus (HIV), maturation of released particles into the infectious form requires protease cleavage of Gag and Gag-Pol as well as major rearrangement of the internal virion structure; heterogeneity in maturation state is a source of within-population diversity (3, 4). Virion–virion aggregation adds another layer of heterogeneity to virus populations. Aggregates may attach, enter, and infect as units, potentially altering infection dynamics and facilitating genetic interactions among co-delivered viral genomes (5, 6). Essentially, no two virus particles in a population are the same. Therefore, single-particle methods are needed to characterize within-population diversity that traditional bulk assays cannot resolve.

The most common methods for detecting particle-level structural heterogeneity in virus populations are electron microscopy (EM) and optical microscopy, which support quantitative comparisons among individual virions. However, sample preparation often required for EM, such as purification, staining, or sectioning, may alter particle morphology or bias the representation of subpopulation abundance relative to the original sample (7, 8). Single-virion optical microscopy, such as total internal reflection fluorescence (TIRF) microscopy, can provide dynamic information on individual particles at lower sample concentrations but requires immobilization before imaging (9). Modern super-resolution techniques can provide information about morphology (10) and subviral composition (11), but remain relatively specialized, requiring dedicated microscopy equipment and expertise. Most importantly, both electron and optical microscopy are inherently low-throughput, requiring hours to obtain and analyze phenotypic data from a virus sample. Limited sampling depth constrains robust characterization of phenotypic distributions, particularly in low-yield samples or for rare subpopulations. Flow virometry addresses many of these limitations by enabling rapid analysis of individual virus particles present at low concentrations in solution with minimal sample processing, using flow cytometers that are often accessible to virology laboratories (12, 13). Physical properties of particles are assessed through 405-nm light scatter, while binding of fluorescent antibodies to abundant surface antigens enables specific detection of virions above background. Virus samples can be analyzed without prior purification, with large numbers of individual virions sampled directly in suspension.

The first documented use of flow virometry followed shortly after the introduction of flow cytometry: in 1979, the technique was used to detect T2 bacteriophage and reovirus top component by light scatter (14). Subsequent applications of flow cytometry to virus particles expanded to detecting and sorting virus subpopulations, studying virion assembly and maturation, and isolating individual virus particles for genomic or infectivity analysis (12, 15, 16). Flow virometry has seen relatively broad application in HIV research, including characterization of heterogeneity in virion-associated envelope glycoprotein (Env) and differential incorporation of host proteins in viral envelopes (17, 18). Furthermore, fluorescent HIV-1 particles have also been isolated by flow-based sorting and retained infectivity (19). More recently, we employed flow virometry to measure the size and shape of IAV virions and demonstrated that their shape can rapidly change in response to environmental conditions (2). Nonetheless, flow virometry remains under-utilized outside of the HIV field, likely due to the lack of an easy-to-adopt protocol for use with different virus families and experiment goals.

Here, we detail a simple and adaptable workflow for applying flow virometry across diverse virus systems and demonstrate its utility for assessing multiple dimensions of viral physical diversity. We first establish practical labeling and acquisition conditions for robust detection and counting of particles from multiple IAV strains and diverse enveloped and non-enveloped viruses. We next apply flow virometry to detect virion interactions at different scales, including antibody binding to individual virions and virion–virion aggregation. We then illustrate how flow virometry can provide a physical-particle denominator for orthogonal bulk measurements by combining it with droplet digital polymerase chain reaction (ddPCR) to quantify viral population ploidy and defective viral genome content (20). We further illustrate how flow virometry can measure dynamic processes in real time through analysis of lipid-mixing kinetics during IAV membrane fusion (21). Finally, we describe the use of flow-based sorting to separate viral subpopulations based on virion morphology while retaining their infectivity. Together, these applications establish flow virometry as an adaptable approach for studying within-population viral diversity and dynamics, with potential utility for other nanoscale systems such as extracellular vesicles, subcellular organelles, large protein complexes, and lipid nanoparticles.

## Results

### Flow virometry – an example IAV workflow and general signal-to-noise considerations

In flow virometry, 405-nm (violet) side scatter (V-SSC) enables detection and counting of nanoscale particles. Scattering intensity is related to the physical features of particles, such as diameter, refractive-index contrast with the surrounding medium, and, for pleomorphic viruses, their shape and orientation (22, 23). We previously applied these characteristics to count and quantify the fraction of filamentous particles in IAV populations across four strains, including in low-yield, attenuated infections (2). Success depended on separating the virion signal from the intrinsic background noise present in sheath or buffer fluids, including contaminating small particles or air bubbles as well as optical and electronic artifacts that fall within the same V-SSC range as virions. This was achieved by using high-affinity, brightly labeled fluorescent antibodies targeting accessible surface epitopes on the most abundant IAV surface glycoprotein, hemagglutinin (HA).

In this section, we present a rapid and robust IAV flow virometry strategy employing fluorescent labeling and outline important considerations for improving signal-to-noise in virion detection. For detecting H1N1 IAV Puerto Rico/8/1934 (PR8), we used high-affinity antibody H36-26 against the surface-accessible HA1 Sb epitope (24) (**Fig. 1A**). 8 x 10^7^ virions/µL were incubated with 20 nM DyLight 550 (DL550)-labeled H36-26 antibody for 30 min at room temperature to allow antibody binding. To minimize background fluorescence from unbound antibody, the virion–antibody mixture was diluted 250-fold into antibody-free buffer without a washing step before analysis by flow virometry. Compared with the unlabeled sample (**Fig. 1A**, *middle*), which partially overlapped with intrinsic background noise (**Fig. 1A**, *left*), antibody labeling clearly separated virion-associated events from noise (**Fig. 1A**, *right*). The labeled virions resolved into at least two partially overlapping subpopulations, corresponding to spherical virions and filamentous virions and/or aggregates (2, 20).

**Figure 1.**
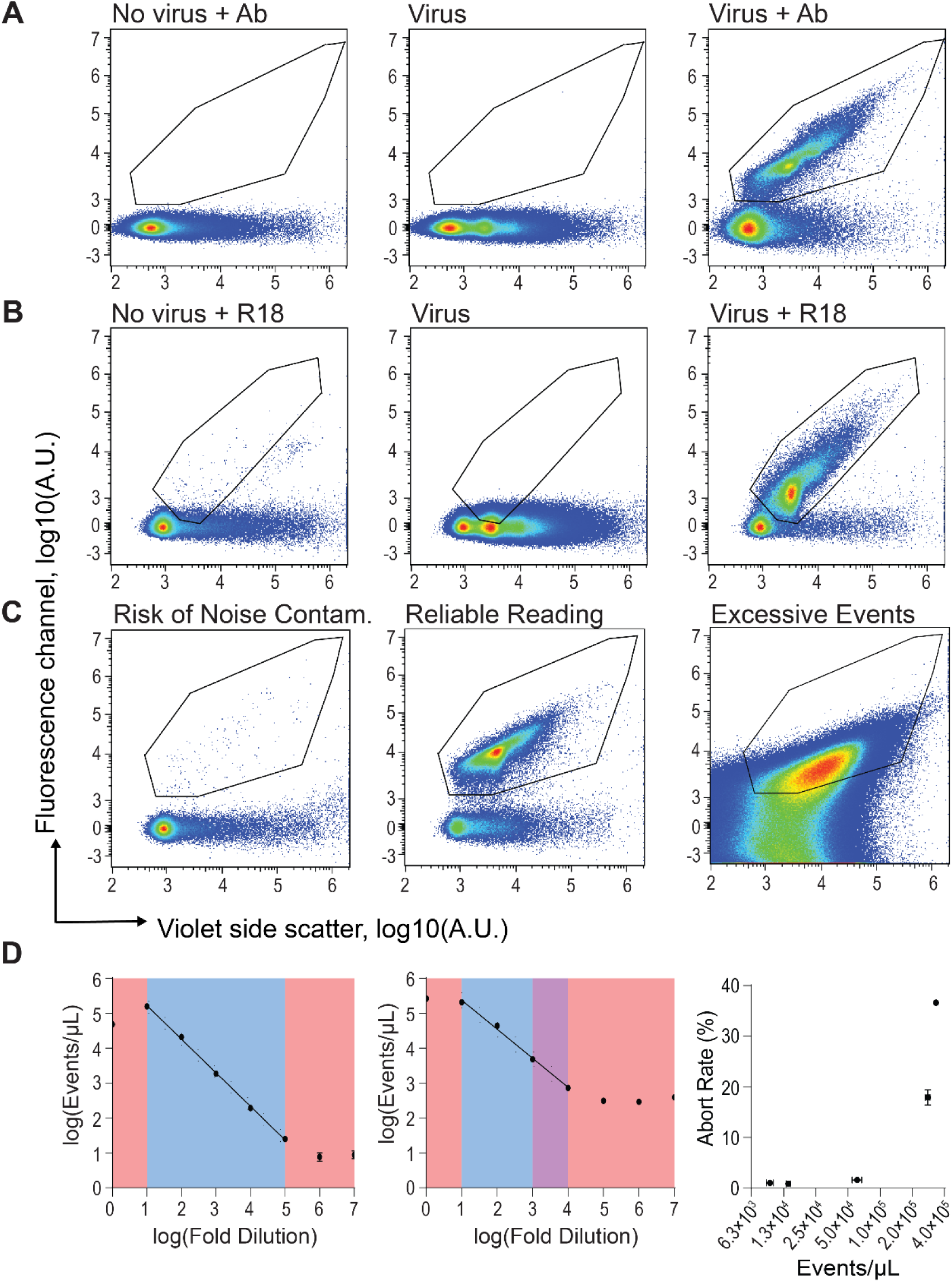
Fluorescent labeling supports accurate virion counting across a wide range of particle inputs. (A) *Flow virometry with antibody labeling*. Events were triggered on V-SSC area with a cutoff of 800. *Left*, a no-virus control showing background noise from buffer alone; *middle*, PR8 virus stock without antibody staining; *right*, PR8 virus stock stained with 20 nM DL550-conjugated antibody H36-26. (B) *Flow virometry with lipophilic dye labeling*. *Left*, a no-virus, 10 μM R18-only control; *middle*, PR8 virus stock without staining; *right*, PR8 virus stock stained with 10 µM R18. Free dye was removed from labeled virions using a gel-filtration PD-10 column. The no-virus, R18-only control was prepared analogously except that virus was omitted. (C–D) *Defining the accurate counting regime in flow virometry*. (C) Representative antibody-stained samples showing, from left to right, low virion event density with a risk of noise contamination, an optimal event density for reliable acquisition, and excessive event density causing unreliable measurements due to incomplete labeling and signal distortion (see also panel D for increased abort rates). (D) Serial dilution analysis defining the linear counting range for stained samples (*left*) and unstained samples (*middle*), and abort rate as a function of event concentration (*right*). Red-shaded regions show virion concentration regimes with unreliable measurements due to low or excessive event density. Purple-shaded region denotes a ‘caution zone’ where noise contamination may be unacceptably high depending on accuracy needs. Blue-shaded regions indicate an ideal measurement range.

In the absence of a suitable antibody, particularly for emerging viruses lacking specific reagents, antibody-independent labeling methods may enable sufficient separation from noise. These include direct conjugation of viral proteins to NHS ester dyes or incorporation of lipophilic dyes into viral membranes; both strategies require purified virus preparations. We illustrate this approach using R18 labeling of PR8 IAV (**Fig. 1B**). Purified PR8 at 1 x 10^8^ virions/µL was incubated with 10 µM R18 at room temperature for 1 h, followed by removal of unincorporated dye using a PD-10 desalting column. This cleanup step was necessary to achieve substantial separation of virion fluorescence from background, although separation was less complete than with antibody labeling (**Fig. 1B**, *right*). The no-virus control contained some background events within the virus particle gate, presumably representing dye aggregates or other supramolecular assemblies that were not removed by the PD-10 column (**Fig. 1B**, *left*). The requirements for virus purification and dye removal represent key limitations of nonspecific labeling approaches, as such manipulations can lead to particle loss and/or aggregation.

The success of flow virometry depends on minimizing background noise and optimizing virion fluorescence. The goal is to achieve a low background event rate within the particle gate for unlabeled measurements and to prevent background noise from overlapping with virion fluorescence signals for labeled measurements. The maintenance and running conditions should be established empirically for each instrument and adjusted as needed. In addition to following the manufacturer’s maintenance instructions, we run a buffer well for a total of 6 min at 10 µL/min before sample acquisition to flush the fluidic line. Furthermore, because the noise level is often elevated at the beginning of runs before the flow rate stabilizes, data from the first 90 s of sample acquisition are typically excluded from analysis (**Fig. S1B**). We also recommend acquiring samples at a slow flow rate, typically 10–20 µL/min, to reduce noise. After each sample, we perform a 20 s backflush to rinse the sample line and clean the sampler tip. After every six samples, a 2 min run at 10 µL/min with buffer only is performed to periodically flush the line (**Fig. S1C**).

Optimization of virion fluorescence depends on both targeting abundant, exposed virion components (e.g., proteins or lipids) and using high-affinity antibodies efficiently labeled with a bright fluorophore. For IAV, HA is an especially effective target because it is the most abundant and accessible virion surface protein. Small, spherical virions display approximately 300–400 HA trimers, but only about 50 NA and 10 M2 tetramers (25, 26). Consistent with this principle, antibodies targeting the HA head domain generally produce stronger virion labeling than antibodies targeting the less accessible HA stem region. High-affinity antibodies are also important, ideally achieving saturating binding at low-nanomolar concentrations and exhibiting slow dissociation rates that allow them to remain virion-bound after the dilution before acquisition. In our workflow, antibodies were labeled at an approximately 100:1 molar ratio of NHS-ester fluorophore to antibody; however, fluorophore incorporation varied across antibodies. We recommend multiple antibodies and labeling ratios be compared to empirically select those with the highest labeling efficiency. Fluorophore choice also affected detection, with brighter dyes such as DyLight 550 NHS ester (Thermofisher, Cat# 62263) performing well in our hands and some longer-wavelength fluorophores such as DyLight 650 NHS ester (Thermofisher, Cat# 62266) producing weaker separation from background. Together, these parameters define a practical framework for maximizing virion fluorescence and separating labeled particles from background noise.

### Virion counting by flow virometry

We previously validated the accuracy of particle counting by comparing particle concentrations derived from flow virometry with hemagglutination (HAU) titer across different preparations of the same IAV strain, X31HA/Udorn/1972 (XUd), and found a consistent ratio of 1 HAU to 8.9 x 10^5^ virions (2). Importantly, hemagglutination titer depends on virus–receptor avidity (27) and its relationship to physical virion counts can thus vary among strains. Flow virometry-derived particle concentrations are independent of virion functions such as receptor binding, facilitating comparison across strains. Here we present a practical guide for ensuring accurate counting by flow virometry without relying on an independent validation method.

Accurate virion concentration measurement is possible within a range that permits saturating antibody binding and sensitive particle detection above background after dilution (**Fig. 1C, D**). To determine an input particle concentration range in which virometry yields accurate counts, a dilution series of a labeled, concentrated sample can be analyzed and the data plotted as log_10_(fold dilution) versus log_10_(events/µL) (**Fig. 1D**). Generally, a 250-fold dilution after labeling with 20 nM fluorescent antibody provides good separation of IAV virion fluorescence from buffer background, so even the highest particle concentration shown represents the concentration after this initial dilution. Deviation from a straight line with the slope of negative one indicates inaccurate counting, which can result from insufficient signal above background at low concentration and an excessively high event rate at high concentration. In the example shown, the lowest virion concentration allowing accurate particle counting above background noise was ∼10–20 virions/µL (**Fig. 1D**, *left*), which corresponds to a minimum virion concentration during labeling of ∼2,500–5,000 virions/µL. While we have achieved up to fivefold greater sensitivity in past measurements (2), this performance depends on meticulous instrument maintenance and alignment. The lower concentration limit is even more restrictive for unlabeled samples, where V-SSC noise significantly overlaps the virion signal (**Fig. 1D**, *middle*). Importantly, label-free flow virometry cannot distinguish increases in background that may accompany serial runs from virion signal, and therefore extra care must be taken to minimize noise before each sample run. In all cases, counting accuracy at low virion concentrations can be improved by pretreating plates with 0.1% BSA, followed by thorough rinsing and drying before sample addition; blocking the plastic surfaces prevents virion loss due to adsorption (**Fig. S2**). At the other end of the concentration range, a particle concentration that is too high might exceed the labeling capacity of the available antibody, reducing virion fluorescence and causing a loss of separation from background noise (**Fig. 1C**, *right*). Even when antibody concentration is sufficient, highly concentrated samples need to be diluted before acquisition. We measure an upper limit of ∼10^5^ virions/µL, above which the abort rate exceeds the acceptable limit of ∼5% (**Fig. 1D**, *right*). Aborted events occur when multiple virions pass through the detector simultaneously and are not counted as virion events. Consistent with the high abort rates, particle counts deviate from the trend line at concentrations exceeding ∼10^5^ virions/µL (**Fig. 1D**, *left*). Together, these limits establish that raw input particle concentrations for direct measurement with flow virometry can span five orders of magnitude while maintaining high-quality quantification (**Fig. 1D**, *left*).

### Universal IAV labeling strategy for flow virometry

We have previously employed flow virometry to characterize particle shape diversity in IAV populations (2). Using four IAV strains belonging to two subtypes and spanning ∼80 years of antigenic evolution, we showed that particle size distributions correlated with infection efficiency, which was defined as the progeny produced per input virion. Under this model, a greater prevalence of spherical virions results from more efficient infection, leading to the prediction that a predominantly spherical morphology may indicate greater adaptation of an IAV strain to a given host cell (2, 28). To directly test this hypothesis, a labeling strategy broadly applicable across IAV strains and subtypes is needed.

To develop a subtype-independent labeling reagent for IAV, we designed a single-chain version of the broadly cross-reactive antibody MEDI8852 (scMEDI), which recognizes an HA stem epitope conserved across IAV subtypes (29). To increase scMEDI stability and provide sufficient fluorescent signal, we fused it to monomeric superfolder GFP (scMEDI-moxGFP) (20). The small size of the single-chain fragment limited the brightness achievable with NHS ester labeling, while direct fluorophore conjugation was expected to further destabilize the already unstable single-chain construct. The size of scMEDI-moxGFP nonetheless remained small (i.e., comparable to that of a Fab fragment), providing better accessibility to its HA base epitope than full-length MEDI8852 IgG (30). The superfolder GFP derivative enabled high expression while the monomerizing V206K mutation prevented virion aggregation mediated by GFP–GFP interactions during labeling.

Using scMEDI-moxGFP, we successfully labeled IAV strains spanning the H1, H3, and H5 subtypes, demonstrating its broad applicability for flow virometry of diverse IAV strains (**Fig. 2A**). Clear separation of virion fluorescence from noise was evident across IAV samples, and, consistent with our previous flow virometry using HA head-binding antibodies (2), the scatter plots revealed heterogeneity in virion signal. The plots contained a dominant population composed of small, spherical particles accompanied by events extending to higher fluorescence and V-SSC values, representing filamentous particles, virion aggregates, or both (2, 20). Notably, these patterns revealed differences in the larger virions among strains: in A/CA/07/09(H1N1), they emerged as a ‘shoulder’ continuous with the spherical population; in A/Perth/16/09(H3N2), they were the least abundant; and in A/TX/37/24(H5N1), they displayed a ladder-like appearance distinct from the spherical virion peak. With scMEDI-moxGFP, we establish an approach for rapidly assessing the particle size distributions of circulating and emerging IAV isolates with potential application as a screen for their adaptation to different host cells.

**Figure 2.**
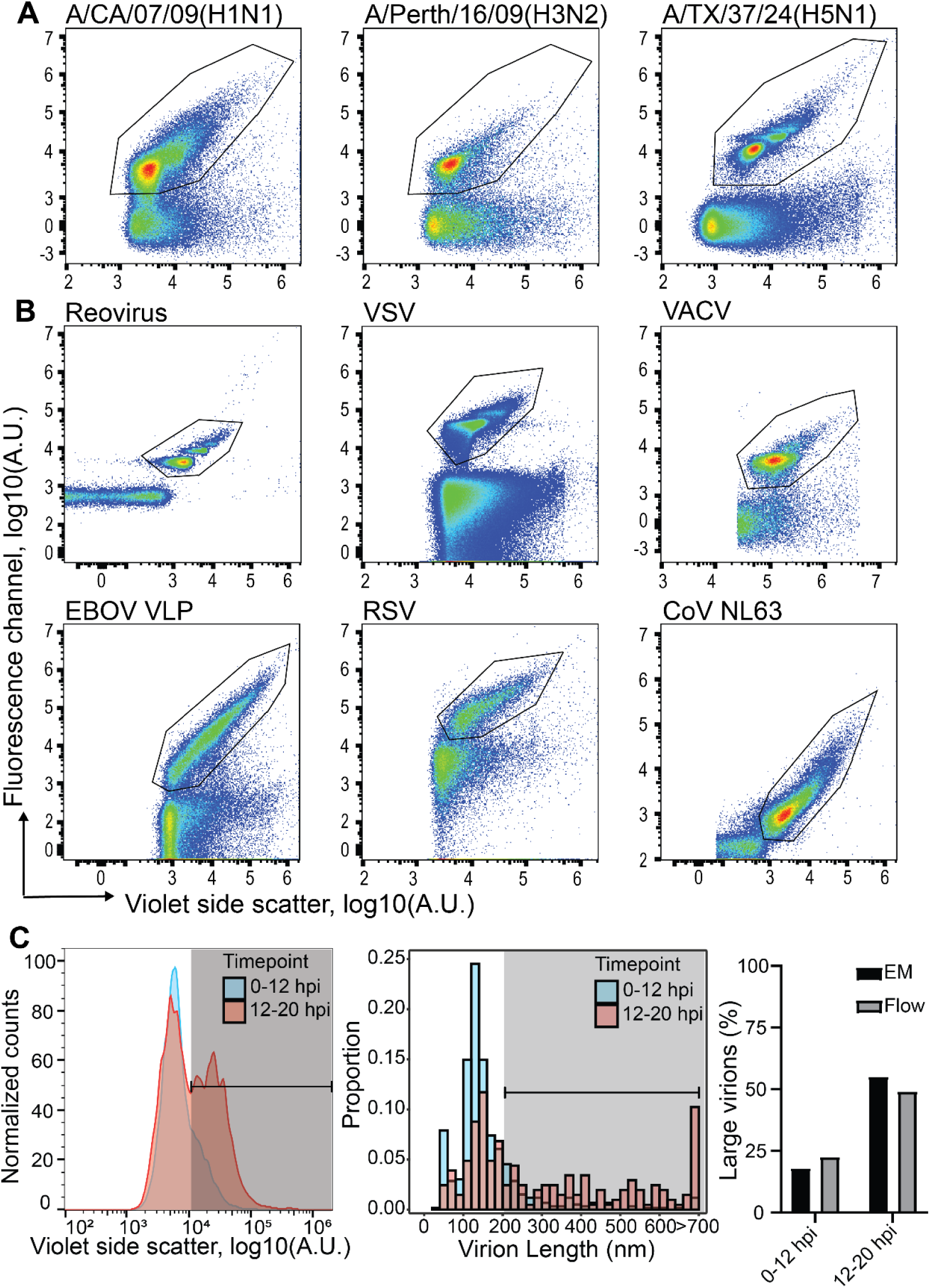
Flow virometry with fluorescent virion labeling is broadly applicable across diverse virus species. (A) *Broad detection strategy across IAV subtypes*. A/California/07/2009 (H1N1), A/Perth/16/2009 (H3N2), and A/Texas/37/2024 (H5N1) stained with scMEDI-moxGFP. (B) *Flow virometry gallery across diverse virus species*. Mammalian orthoreovirus type 1 Lang (reovirus) stained with a single-chain fragment of the 10C1 antibody fused to monomeric superfolder GFP (10C1 scFv-moxGFP); vesicular stomatitis virus (VSV, Indiana serotype) stained with DyLight 488-8G5F11 IgG; vaccinia virus (VACV, WR (tk-), extracellular enveloped virus) stained with LA5 DyLight 488-IgG; Ebola virus-like particles (EBOV VLPs) incorporating GFP-fused matrix protein VP40; respiratory syncytial virus (RSV) strain A2 stained with DyLight 488-D25 IgG; human coronavirus NL63 (CoV-NL63) stained with the lipophilic dye R18. (C) *Correlation of filamentous particles detected by flow virometry (left) and by EM (right).* XUd was harvested from MDCK-Siat1 cells infected at MOI 6 and measured by both flow virometry and EM at ≤12 h (*blue*) and 12-20 h (*red*). *Left, temporal changes in XUd virion size revealed by flow virometry*. V-SSC profiles of XUd; the filamentous-virion gate is indicated by the black line and gray shading. *Middle*, *temporal changes in XUd virion lengths revealed by EM*. Histogram of virion length distributions; ≤12 h (n = 619 virions) and 12–20 h (n = 231 virions) from 66 and 107 micrographs, respectively. Virion lengths exceeding the 200-nm cutoff are indicated by the black line and gray-shaded area. *Right*, *comparison between virion-size measurements by EM and flow virometry*. EM (*black bars*) and flow virometry (*gray bars*) data for the two collection intervals.

### Flow virometry as a generalizable approach – single-particle analysis of viral pleomorphy

While morphological heterogeneity is well recognized for pleomorphic enveloped viruses such as IAV, particle-level heterogeneity may be less apparent in viruses with more constrained architectures. For such viruses, genetic diversity within populations is well recognized, yet virions are often still conceptualized as structurally uniform copies of one another. In fact, individual particles often differ in morphology, size, maturation state, molecular composition, and aggregation state. These particle-level differences may affect infectivity, stability, immune recognition, and transmission, yet they are difficult to resolve with bulk assays. Flow virometry can enable direct single-particle analysis of these features, revealing their underlying distributions. In this section, we describe an extension of flow virometry to a panel of additional viruses, including enveloped and nonenveloped forms and symmetric and pleomorphic virion architectures.

We selected a panel of enveloped and non-enveloped viruses that varied in morphology, structural uniformity, and particle size (**Fig. 2B**). We included reovirus as a representative non-enveloped virus with a uniform icosahedral structure and a diameter of approximately 80 nm. We labeled it with antibody 10C1 (31), which targets σ3, an outer capsid protein present at 600 copies per fully matured virion (32). As expected, the majority of reovirus virions formed a compact cluster in the V-SSC versus fluorescence scatter plot (**Fig. 2B**, *top left*). Notably, minor but distinct subpopulations displayed higher V-SSC and fluorescence signals. Given the defined icosahedral structure of reovirus, these subpopulations likely represent doublets, triplets, or higher-order aggregates rather than larger single virions. We also analyzed vesicular stomatitis virus (VSV) and vaccinia virus (VACV), two enveloped viruses with distinct but relatively uniform morphologies. VSV virions are bullet-shaped, approximately 70 × 200 nm (33), whereas VACV virions are brick-shaped with slightly rounded edges, approximately 360 × 270 × 250 nm (34). When stained with antibodies targeting their major surface glycoproteins (VSV G and VACV D8L) and analyzed by flow virometry, both viruses formed single predominant populations in the scatter plots (**Fig. 2B***, top middle and right*). Like reovirus, both displayed minor subpopulations of larger particles that formed a ladder-like pattern. As the VSV stock was propagated under conditions that minimize defective interfering particle production, extensive morphological heterogeneity was not expected. Indeed, few events were observed below the main particle population in either V-SSC or fluorescence, and the larger-particle subpopulations therefore most likely represent successive orders of virion aggregation (**Fig. 2B***, top*). We similarly hypothesize that the larger-particle subpopulations in the VACV sample represent virion aggregates. Together, these experiments extend flow virometry to phylogenetically and structurally distinct viruses and suggest that virion aggregation is more commonplace among viruses than previously appreciated.

We next examined additional pleomorphic viruses using different labeling strategies. We produced fluorescent Ebola virus-like particles (EBOV VLPs) by co-expressing matrix protein VP40 fused to GFP with glycoprotein GP; we labeled respiratory syncytial virus (RSV) virions with D25 IgG, which targets an epitope at the apex of the prefusion F glycoprotein; finally, we labeled human coronavirus NL63 (CoV NL63) virions nonspecifically with the lipid dye R18. Each labeling strategy resulted in efficient separation of virus particle fluorescence from noise, although, as with IAV, separation was less complete for the nonspecific R18 labeling. In each case, flow virometry scatter plots revealed continuous populations with a prominent region of high event density representing smaller virions and subpopulations with higher V-SSC and fluorescence intensities, consistent with larger virions and/or virion aggregates (**Fig. 2B**, *bottom*). These combined results extend our observations in IAV, showing that preparations of diverse viruses are not monodisperse or composed of uniformly shaped virions but instead contain heterogeneous particle subpopulations that may include larger virions, filamentous particles, and/or aggregates.

In structurally uniform viruses, larger particles detected by flow virometry likely represent aggregates; in pleomorphic viruses, however, they may instead reflect larger individual virions. Electron microscopy (EM) can be used to assess the presence of filamentous virions among larger particles detected by flow virometry. We previously found that XUd IAV grown in MDCK-Siat1 cells produced a higher proportion of large particles later in infection (2). To examine the morphological basis of this finding, we compared flow virometry and negative-stain EM measurements from two collection intervals during an XUd infection of MDCK-Siat1 cells at an MOI of 6. Virions were collected during 0–12 and 12–20 h of infection. Flow virometry of virions released during the first 12 hours of infection showed a dominant small virion peak with a minor shoulder extending toward higher V-SSC values (**Fig. 2C**, *left*). This distribution corresponded to the predominantly small virion population observed by EM (**Fig. S3**; **Fig. 2C**, *middle*). Later in infection, the V-SSC distribution shifted toward higher values (**Fig. 2C**, *left*), and EM confirmed an increased proportion of filamentous virions (**Fig. S3**; **Fig. 2C**, *middle*). The proportion of particles within the large-virion gate by flow virometry agreed with the proportion of virions exceeding 200 nm in length by EM (**Fig. 2C**, *right*). Although two matched samples are insufficient to rigorously define the boundaries between small and large virions by either method, the concordant temporal changes measured by EM and flow virometry support V-SSC as a readout of population morphology in this system. These data do not exclude aggregates within the higher-V-SSC population but suggest that filamentous virions are the predominant source of its temporal increase. Thus, flow virometry captures changes in particle size distributions, while EM provides the morphological information needed to unambiguously identify filaments.

### Detection of antibody binding and virion–virion interactions by flow virometry

Physical interactions between virions and binding partners, including other virions, antibodies, or complement proteins, can alter particle properties in ways that are difficult to resolve using conventional bulk assays. Traditional approaches can measure binding or aggregation indirectly but often require ensemble averaging, biochemical separation, or imaging-based workflows that are low throughput and labor-intensive. Flow virometry provides a direct single-particle approach to detect virion interactions by measuring changes in light scatter and/or fluorescence. For sufficiently concentrated virus, unlabeled virions can be detected above background using V-SSC alone (**Fig. 3A**; see also **Fig. 1A)**; we exploited this to measure increase in virion V-SSC signal upon binding of unlabeled antibodies. We incubated purified virions at 10^6^ particles/µL with a panel of eight anti-HA antibodies at 100 nM and measured V-SSC by flow virometry; each of the eight antibodies produced a noticeable shift toward higher V-SSC values (**Fig. 3B**). The magnitude of the shift varied by epitope class, with greater increases in V-SSC for HA head-than HA stem-binding antibodies. This difference may reflect variation in epitope accessibility and, consequently, antibody occupancy. Additionally, an antibody extending from the virion surface and primarily increasing its effective cross-sectional area (a head-binder) may affect V-SSC differently than one binding closer to the virion surface within the pre-existing HA layer and primarily increasing local protein density (a stem-binder). Nonetheless, the epitope-dependent V-SSC shifts reveal that flow virometry can not only detect antibody binding to intact virions but may also distinguish binding to different epitope classes.

**Figure 3.**
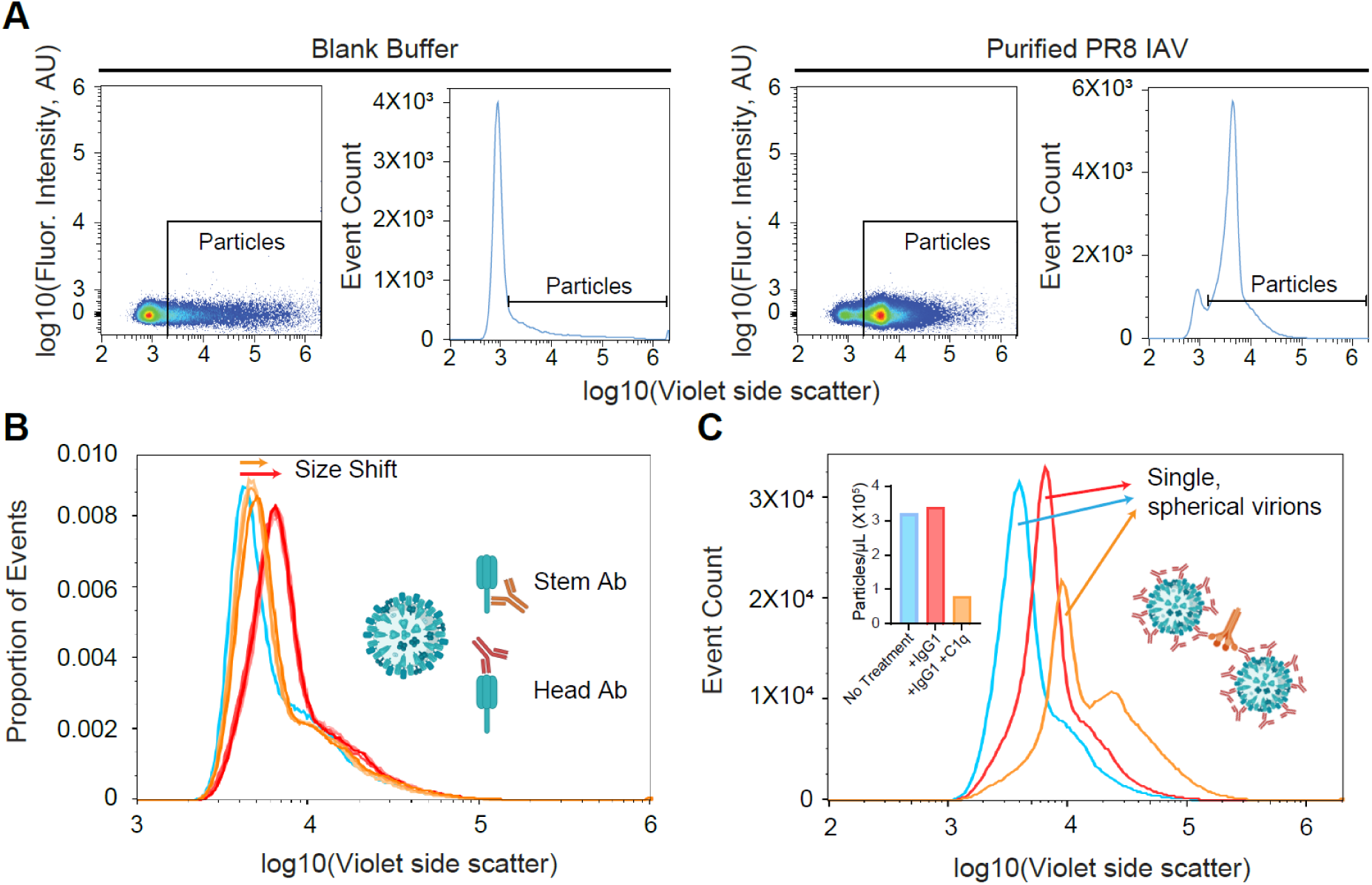
Flow virometry detects interactions across scales, from virion–antibody binding to virion–virion aggregation. (A) V-SSC vs. fluorescence scatter plots and V-SSC histograms for the buffer blank (*left*) and purified PR8 (*right*) with the particle gate indicated. (B) Unlabeled PR8 virions were analyzed either untreated (blue) or after incubation with unlabeled antibodies: HA stem-binders CR9114, MEDI8852, Fi6, or 16G8 (*shades of orange*), or HA head-binders H36-26, H28D14, Y8IA6, or H35C12 (*shades of red*). (C) Unlabeled PR8 virions were analyzed either untreated (*blue*) or after incubation with unlabeled HA head-binding antibody H28D14 alone (*red*) or together with C1q (*orange*).

Antibody-induced V-SSC shifts can also aid in screening candidate antibodies for flow virometry. Measuring this response across a range of antibody concentrations can identify candidates that achieve maximal V-SSC shifts at low nanomolar concentrations without requiring fluorescent labeling of each antibody. Suitable antibodies should also have low dissociation rates, which can be evaluated by monitoring decay of the antibody-induced V-SSC signal after dilution. Candidates with high affinity and low off-rates can then be conjugated to NHS ester dyes and tested for retention of virion-binding activity and sufficient fluorescence to separate labeled virions from background. Antibodies with high affinity, slow dissociation, preserved binding after dye conjugation, and complete fluorescence separation from background are optimal flow virometry reagents.

In addition to detecting protein–virion interactions, flow virometry can also quantify treatment-induced aggregation. We incubated virions with an HA head antibody with or without the Fc-binding protein C1q and analyzed the samples by flow virometry (**Fig. 3C**). We used 70 µg/mL C1q to approximate its physiological concentration in human serum (35). C1q caused both a reduction in the overall particle concentration and an increase in the proportion of larger aggregated particles. Based on these changes, we estimate an average of three virions per aggregate. Aggregation did not affect all particles uniformly, as a subpopulation remained monodisperse, underscoring an advantage of single-virion analysis for revealing nonuniform effects that are obscured by bulk measurements. Together, these results illustrate the versatility of flow virometry for assessing interactions across scales, from protein–virion binding to virion–virion interactions.

### Flow virometry enables particle-normalized quantification of viral population properties

Virus populations contain multiple layers of heterogeneity that are difficult to capture using any single measurement. By combining flow virometry with orthogonal quantitative assays, bulk molecular measurements can be normalized to particle counts or another flow virometry-measured property, such as HA abundance per virion. This creates a general framework for using bulk measurements of genome content, protein abundance, lipid content, enzymatic activity, or infectious units to quantify average molecular or functional properties per particle or to relate them to other particle characteristics.

This framework can be readily applied to measure viral population ploidy, defined as the average number of genome copies per particle. Across populations, ploidy can vary due to factors such as virion aggregation, alterations in genome packaging, clustering of virions within vesicles, or attachment to bacterial cells (5, 36–39). As a fundamental biological trait, viral population ploidy may influence evolutionary outcomes by affecting processes such as reassortment in segmented viruses; in IAV, reassortment underlies the emergence of pandemic strains (40). We previously combined particle concentration measurements by flow virometry with orthogonal measurements of genome copy number by droplet digital PCR (ddPCR) from the same virus sample (20). This enabled us to derive viral population ploidy even when virus particle production was reduced by antibody treatment or the NA inhibitor (NAI) oseltamivir. Oseltamivir promotes retention of virion–virion contacts during budding by inhibiting NA-mediated cleavage of sialic acid on virions (20, 41).

To demonstrate this principle, we performed a similar experiment here, inducing an increase in viral population ploidy with oseltamivir and quantifying it by combining ddPCR with flow virometry as follows (**Fig. 4**). To quantify genome copy number, we performed viral RNA extraction, reverse transcription, and ddPCR (**Fig. 4A**). To minimize the likelihood of a single RNA template being replicated multiple times, we used Moloney murine leukemia virus reverse transcriptase (M-MuLV RT), which possesses RNase H activity that degrades the RNA template during reverse transcription. To correct for potential gain or loss during sample processing, we included a known number of VSV particles, also quantified by flow virometry (**Fig. 2B**), as an internal control at the RNA extraction step. For this calculation, we assumed that each VSV virion contains one genome. This assumption is supported by VSV’s strict requirement for genome packaging during virion assembly and the dependence of virion size on the length of the packaged genome, suggesting that empty particles are rare in the population. We note that this is an imperfect assumption given that the VSV stock likely included a small subpopulation of virion aggregates (**Fig. 2B**). Nonetheless, we used the raw VSV genome-to-particle ratio as a calibration coefficient to adjust IAV measurements, approximating absolute genome-to-particle ratios. As defective viral genomes (DVGs), characterized by large internal deletions, are common among RNA viruses, we additionally designed primers to enable quantification of DVGs alongside the full-length IAV PB2 and HA segments (**Fig. 4B**). To quantify total segment copies (full-length and truncated segments), we targeted the primers to terminal regions of each segment that are typically retained in DVGs. To specifically quantify full-length segments, primers targeted an internal region which DVGs usually lack (42). In each case, the primers were designed to generate ∼200-nt amplicons.

**Figure 4.**
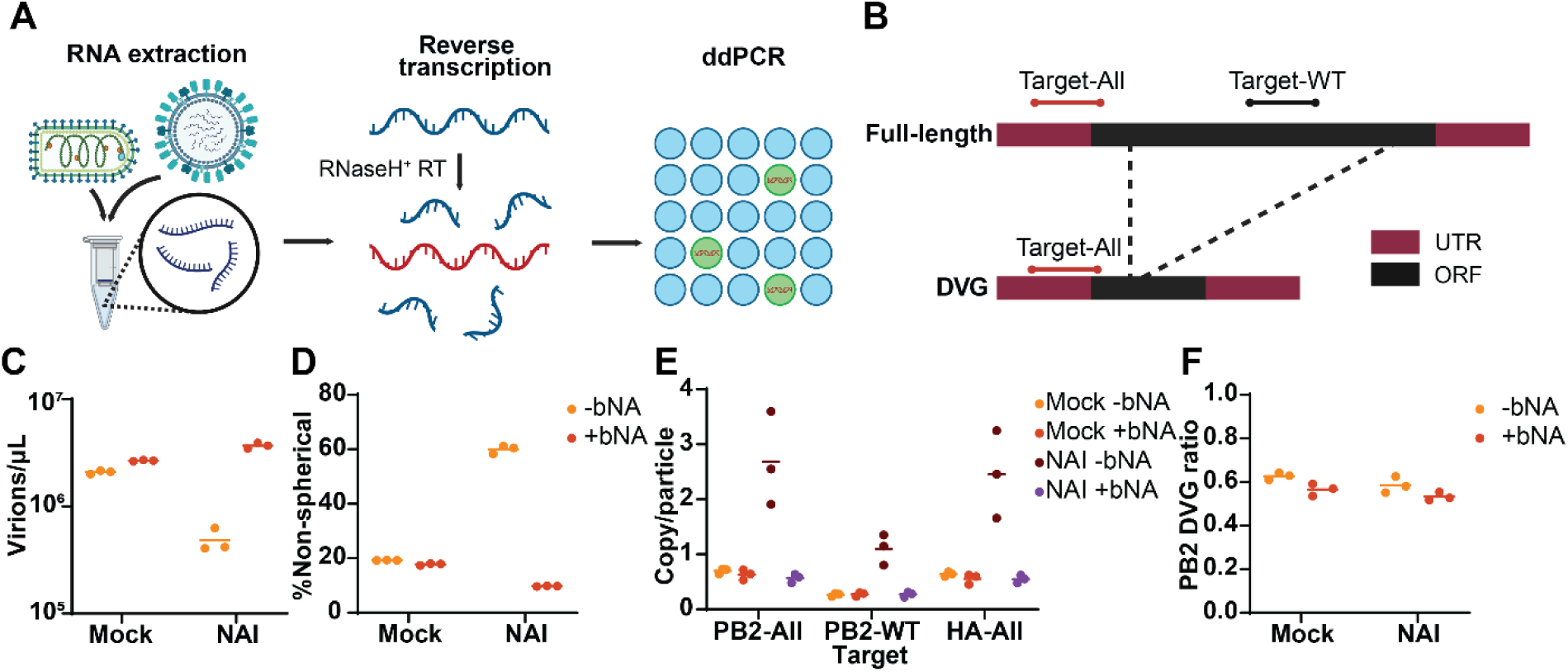
Flow virometry combined with ddPCR enables quantification of IAV population ploidy and DVG abundance. (A) *Scheme for quantifying viral genomes by ddPCR*. Known numbers of VSV and IAV virions are combined and subjected to RNA extraction, reverse transcription with RNase H^+^ reverse transcriptase, and ddPCR analysis. (B) *Primer design for distinguishing full-length and truncated segments by ddPCR*. Terminal regions of each segment are retained in truncated segments, whereas internal sequences are frequently deleted. (C–F) *Proof of principle for measuring IAV population ploidy and DVG abundance*. MDCK-Siat1 cells were infected at MOI 0.6 IU/cell. Bacterial NA (bNA) was added to the cells at 4 hpi to remove sialic acids from the cell surface. 1000 nM oseltamivir was added at 22 hpi to inhibit viral NA activity post-entry. Virions released into the supernatant at 24 hpi were analyzed for particle concentration (C), percentage of non-spherical particles (D), genome-to-particle ratios for the PB2 and HA segments (E), and DVG fraction for the PB2 segment (F). The PB2 segment was chosen because of its known propensity to acquire deletions that generate DVGs.

We demonstrated the feasibility of combining ddPCR and flow virometry to quantify the ploidy of IAV populations released from infected cells. We treated IAV-infected MDCK-Siat1 cells with bacterial neuraminidase (bNA) at 4 h postinfection (hpi) to facilitate cleavage of cell-surface sialic acids. At 22 hpi, we replaced the medium with oseltamivir-containing medium and harvested the supernatants at 24 hpi. We quantified released particle counts, the proportion of large particles, and the genome-to-particle ratio. NAI treatment reduced particle counts and increased the proportion of large particles, consistent with virion aggregation (**Fig. 4C, D**). We measured average genome-to-particle ratios of 0.698 for the PB2 segment and 0.627 for the HA segment in mock-treated samples without bNA (**Fig. 4E**), in agreement with previous measurements (43, 44). Consistent with the propensity of PB2 to generate DVGs, DVGs constituted around 60% of PB2 segments across treatments (**Fig. 4F**). Notably, samples treated with NAI exhibited higher genome-to-particle ratios than controls, with an average genome-to-particle ratio of 2.69 for the PB2 segment and 2.46 for the HA segment, but this trend was reversed in the presence of bNA (**Fig. 4E**). These results indicate that NA activity, whether viral or bacterial, inhibits sialic acid-mediated aggregation of IAV. Overall, combining flow virometry and ddPCR enables quantification of viral population ploidy and illustrates a broader framework for integrating bulk measurements with flow virometry-measured particle features.

### Measuring lipid-mixing dynamics at single-virion resolution

Typical sampling rates in flow virometry are on the order of 1,000 to 15,000 events/s, enabling rapid characterization of population-level distributions with high statistical power. This high-throughput sampling can be applied to measure the kinetics of time-varying virus-mediated phenomena. We previously exploited this capability to measure viral lipid-mixing dynamics using an approach we termed single-pair Cytometric Analysis of Lipid Mixing (spCALM) (21). Here, we performed a similar experiment to outline the workflow. R18-fluorescence dequenching is well established as a probe for lipid mixing; when incorporated into a viral membrane at a self-quenching concentration, lipid mixing between viral and target membranes dilutes R18, resulting in dequenching and increased fluorescence intensity. Since the fluorescence of single R18-labeled virions can be resolved by flow virometry (see **Figs. 1B, 2B**), we applied the R18 dequenching principle to a flow virometry-based IAV lipid-mixing assay as follows. R18-labeled virions were bound to liposomes displaying IAV receptors at a low virion-to-liposome ratio to favor binding of at most a single virion per target. We identified virion-liposome pairs as the subpopulation of liposomes positive for R18 fluorescence (**Fig. 5A**) and verified single-virion binding by ensuring superimposable fluorescence distributions between liposome-bound and free virions (**Fig. 5B**). The virion-liposome suspension was then rapidly diluted into low-pH buffer to trigger IAV membrane fusion. As virion-target pairs underwent lipid mixing, their R18 fluorescence intensity increased, forming a distinct bright population. The kinetics of lipid mixing were derived from the ratio of bright, lipid-mixed pairs to total pairs as a function of time (**Fig. 5A**), and lipid-mixing efficiency was defined as this ratio at the reaction plateau (**Fig. 5C**). Sample transport from the lipid-mixing reaction tube to the cytometer flow cell causes a measurement delay (∼10 s in **Fig. 5C**); the duration of this delay depends on the flow rate and length of the instrument’s fluidic tubing. When lipid-mixing lag time is sufficiently long relative to this delay, reaction kinetics can be extracted. This approach allows separate analysis of attachment and membrane fusion while providing a substantially higher sampling rate than microscopy-based single-virion membrane fusion assays.

**Figure 5.**
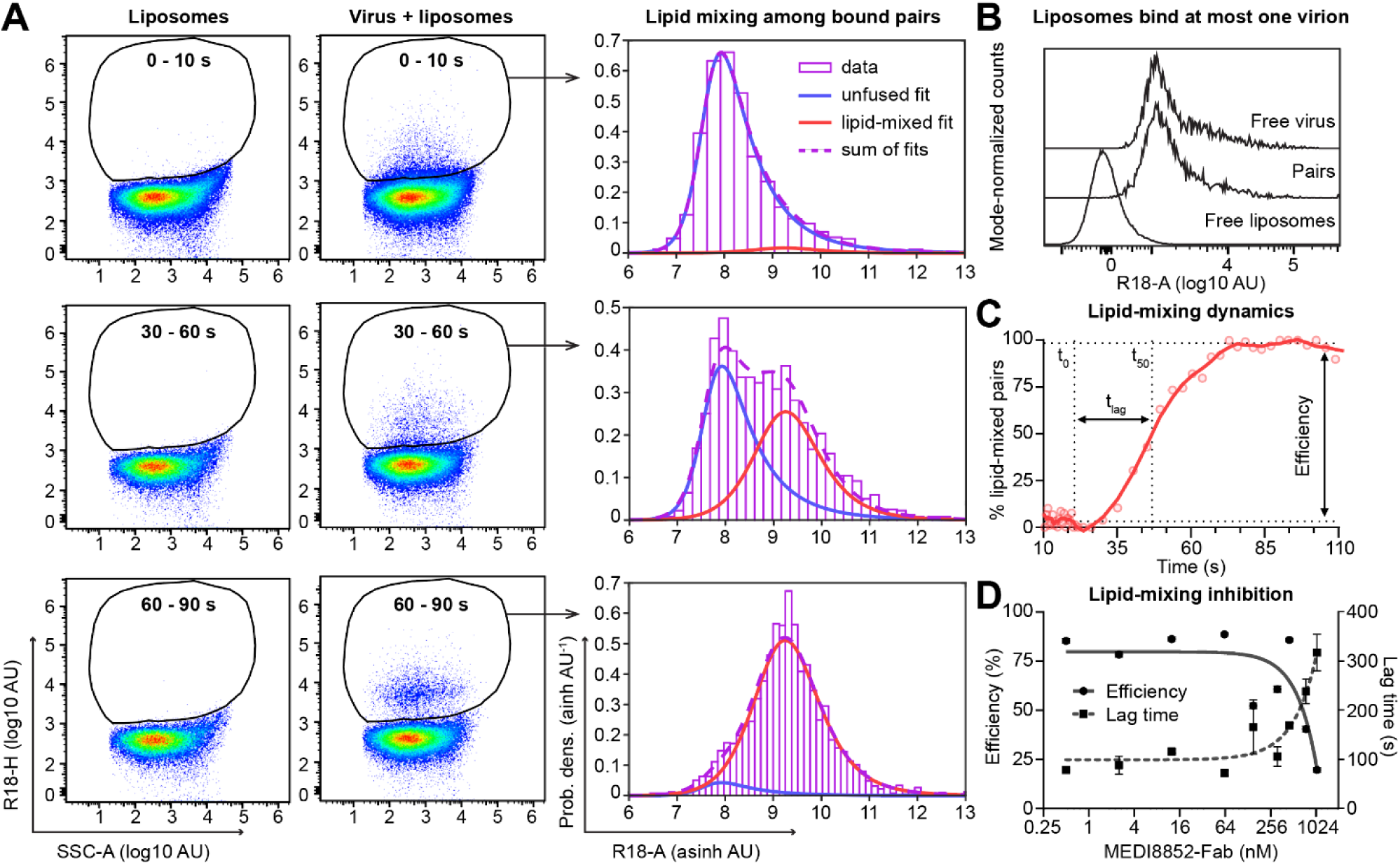
Flow virometry enables real-time analysis of lipid-mixing dynamics and inhibition with single-virion resolution. (A) *spCALM for measuring lipid-mixing kinetics*. Scatter plots of free liposomes (*left*) or liposomes incubated with R18-labeled virions (*middle*) at the indicated time intervals. Virion-bound liposomes (*gate*) are distinguishable from the major population of free liposomes based on fluorescence. pH was lowered at t = 10 s by rapidly diluting the sample into a pH 5.68 buffer. *Right,* the overall R18 fluorescence distribution of bound pairs (*purple bars*) for each time bin is fit as a weighted sum of unfused (*blue*) and lipid-mixed (*red*) reference distributions. The R18 fluorescence intensity of bound pairs increased after low-pH exposure and nearly all pairs underwent lipid mixing by the 60–90 s time bin. (B) *R18 fluorescence distributions of free virions, free liposomes, and virion-liposome pairs.* (C) *Analysis scheme for extracting the efficiency and kinetics of lipid mixing*. We performed the distribution fitting shown in panel A, *right*, for successive time bins, and lipid-mixing efficiency is defined as the percent of lipid-mixed pairs at the reaction plateau. Lag time (t_lag_) is defined as the time interval from the detected pH drop (t_0_) to half-maximal lipid mixing (t_50_). Data are plotted from the time of low-pH dilution (10 s). t_0_ indicates the time when the diluted sample reaches the detector and is greater than 10 s due to the fluidic delay. Bins are chosen to include equal numbers of detected pairs per bin and are more finely spaced than the 30 s intervals in panel A; bin density is highest before t_0_ because acidic buffer addition dilutes the sample. (D) Pretreatment of virions with MEDI-Fab decreases lipid-mixing efficiency and increases lag time in a dose-dependent manner.

spCALM provides a useful platform for probing the mechanism of viral membrane fusion and its inhibition (21). As an example, we treated IAV virions with the Fab fragment of MEDI8852 (MEDI-Fab), which does not interfere with HA-receptor binding but prevents low-pH-induced HA conformational changes required for fusion (1, 21, 29). IAV virions were incubated with a range of MEDI-Fab concentrations, then attached to receptor-bearing liposomes, and fusion was triggered by dilution into pH 5.68 buffer. We observed dose-dependent reduction of lipid-mixing efficiency that coincided with increases in lipid-mixing lag times (**Fig. 5D**). Rapid, high-throughput sampling enabled robust quantification of attenuated lipid mixing under antibody inhibition. Although we describe spCALM here in the context of IAV lipid mixing, its high-throughput kinetic measurements are readily extensible to other dynamic processes, including membrane fusion of other viruses, antibody binding, and viral aggregation.

### Phenotype-based sorting of virion subpopulations

Virus populations are heterogeneous in size, morphology, maturation state, antigen abundance, lipid composition, genome content, and binding to host or immune factors. Isolating subpopulations based on these particle-level features would enable direct testing of how virion phenotypes shape downstream properties such as infectivity, stability, immune recognition, and tropism. The same V-SSC and fluorescence signals used to analyze individual virions by flow virometry can be used to physically sort them if the subpopulations of interest can be resolved by a cell sorter. Sorters generally have lower sensitivity for nanoscale particles than dedicated analytical flow cytometers, in part because sorting requires buffered sheath fluids rather than ultrapure water. As a result, fluorescent antibody binding is required to resolve virions from background on our sorter (Cytoflex SRT). As with flow virometry, the dilution of labeled virus must be empirically determined to achieve optimal sorting efficiency, and collection plasticware should be BSA-treated to minimize particle loss (**Fig. S2**).

To test whether sorter-resolved size differences could support physical separation of virion subpopulations, we developed a sorting protocol to enrich spherical and filamentous subpopulations from a heterogeneous virus stock. For this, we used XUd IAV, for which filamentous virions can be readily distinguished from spherical virions by flow virometry (2) (**Fig. S3, Fig. 2C**). To develop a labeling approach that would not interfere with downstream infection, we engineered a variant of single-chain MEDI8852 fused to enhanced GFP and incorporating a histidine residue in the antigen binding domain (H31 MEDI scFv-eGFP). We hypothesized that eGFP would enable separation of virion signal from noise for sorting, MEDI binding to the HA stem would preserve virion attachment to cells and endocytosis, and H31 protonation in late endosomes would promote antibody dissociation (45), relieving the block on HA conformational changes. Both WT and H31 MEDI scFv-eGFP separated virion fluorescence from noise on the sorter (**Fig. 6A**), and while the WT construct reduced infectivity at the concentrations needed for detection, infectivity of H31-bound virions was unchanged (**Fig. 6B**).

**Figure 6.**
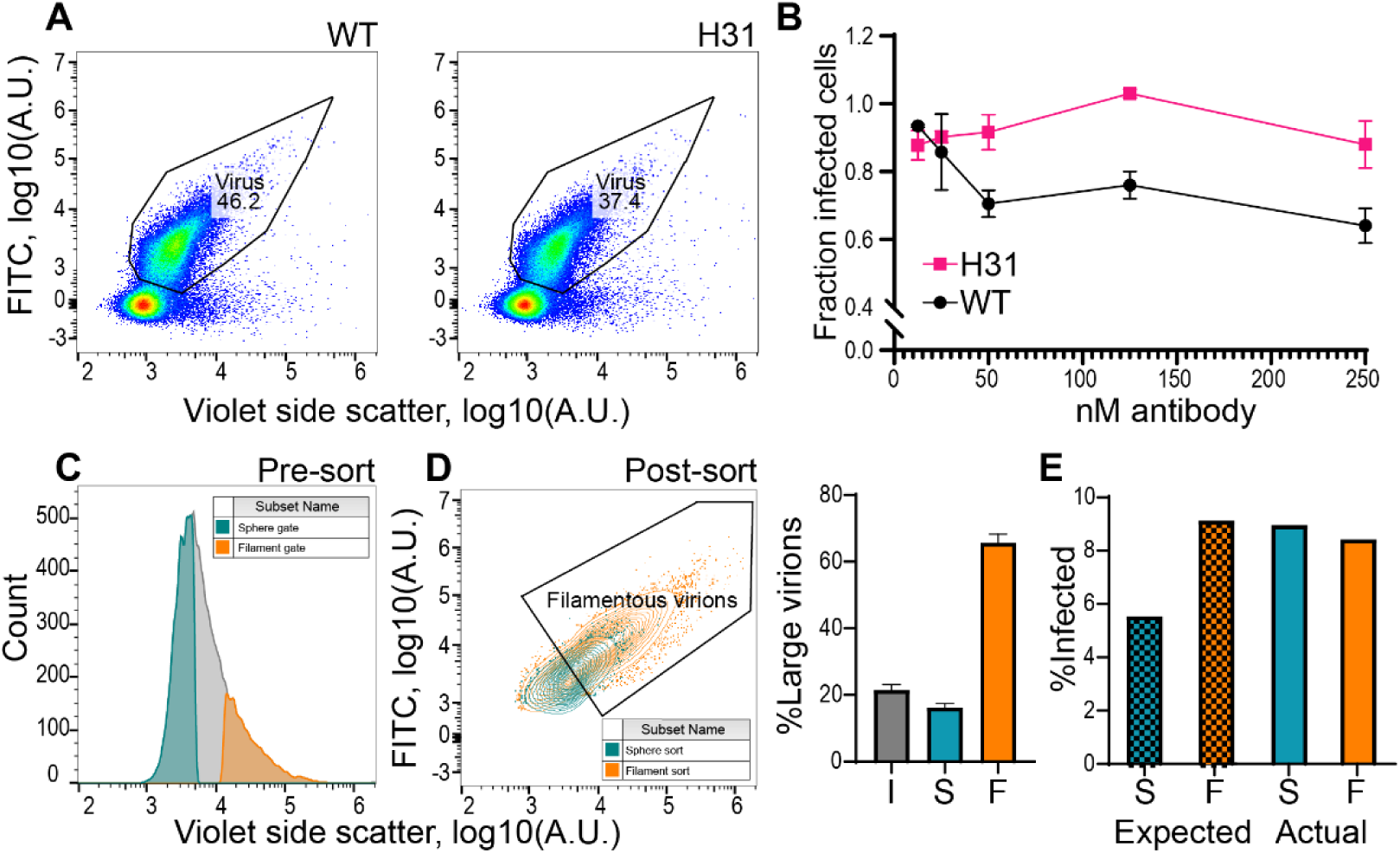
Flow-based sorting enables enrichment of virion subpopulations based on shape. (A-B) *H31 MEDI scFv-eGFP separates XUd virion fluorescence from background on the sorter without reducing infectivity.* (A) V-SSC vs. fluorescence scatter plots acquired on the sorter for XUd virion samples labeled with 250 nM MEDI scFv-eGFP (*left*) or H31 MEDI scFv-eGFP (*right*). (B) XUd infectivity in the presence of increasing concentrations of MEDI scFv-eGFP (*black line, WT*) or H31 MEDI scFv-eGFP (*magenta line, H31*), normalized to an untreated control. (C-E) *Sorting enriches for spherical and filamentous virions while retaining virus infectivity.* (C) V-SSC histogram of XUd labeled with H31 MEDI scFv-eGFP on the sorter, showing the spherical (*teal*) and filamentous (*orange*) virion sort gates. (D) *Flow virometry analysis of sorted virus populations*. Contour plots displaying sorted spherical (*teal*) and filamentous (*orange*) virion populations. Percentage of filamentous virions in sorted populations is shown on the right. I = input, S = sphere sort, F = filament sort. (E) Expected infectivity of sorted samples, calculated from values obtained for the mixed virus population prior to sorting (*Expected*), plotted alongside the measured infectivity of sorted samples (*Actual*). S = sphere sort, F = filament sort.

To sort labeled virus, samples were diluted to 5 × 10^4^ particles/µL. A two-step gating strategy using V-SSC area (V-SSC-A) and V-SSC height (V-SSC-H) was applied to define the virion population (**Fig. S4**), followed by stringent V-SSC-A gates to define spherical and filamentous subpopulations for sorting (**Fig. 6C**). Sorting enriched the spherical population to 82% and the filamentous population to 65%, as determined by subsequent flow virometry analysis (**Fig. 6D**). Notably, sorted virus fractions retained per-particle infectivity on A549 cells comparable to that of the pre-sort bulk population (**Fig. 6E**). Limitations of sorting for downstream applications include substantial sample dilution and low total particle yield, with sorted samples reaching a maximum concentration of 300 virions/µL. Nonetheless, this experiment demonstrates that sorting can enrich virus populations based on size and shape. Using our histidine-engineered H31 MEDI scFv-eGFP label, IAV virions also retained infectivity after sorting.

## Discussion

Flow virometry has emerged as a sensitive, quantitative, and high-throughput method for counting and characterizing single virions within virus populations (2, 14, 18, 19), but widespread adoption has been slow due to the lack of a generalized, accessible protocol (12, 13). We demonstrated its applicability for characterizing within-population diversity across a wide range of enveloped and non-enveloped viruses. We further presented applications including detection of virus–antibody and interparticle interactions, normalization of orthogonal measurements to virion counts, and real-time analysis of dynamic viral processes. Finally, we demonstrated flow-based sorting to separate spherical and filamentous IAV virion subpopulations.

We described several key considerations for successful flow virometry. First, background noise must be minimized because buffer contaminants, debris, microbubbles, and instrument noise can overlap with virion-derived signals; meticulous cytometer maintenance, low flow rates, and routine flushing are therefore important. Second, virion labeling with bright fluorescent probes facilitates separation of virions from background and greatly improves the lower limit of detection, allowing analysis of virions in unpurified infection supernatant and at low virion yields. This is enabled by targeting abundant, accessible virion components with high-affinity binders and bright fluorophores, or by lipid labeling when suitable antibodies are unavailable (**Fig. 1A, B**). Third, particle concentration must remain within a range suitable for accurate counting. At low concentrations, background events constitute an increasing fraction of the measured population, whereas excessive concentrations can compromise labeling and increase coincident detection and abort rates, resulting in inaccurate particle counts (**Fig. 1C**). Finally, the labeling strategy must be adapted to the virus of interest, as illustrated by successful detection of diverse enveloped and non-enveloped viruses using either antibody-based, genetically encoded, or lipid-based labels (**Fig. 2**).

We next illustrated applications of flow virometry that extend beyond basic and static characterization of virus populations to probing virus particle interactions, integrating their counts with orthogonal measurements, and following dynamic changes in populations. Antibody binding to virus particles produces measurable shifts in V-SSC and does not require labeling of either binding partner, allowing native interactions to be measured in solution (**Fig. 3**). The magnitude of the V-SSC shift reflects the extent and/or geometry of binding and can be exploited for rapid screening of suitable flow virometry reagents. V-SSC may provide a broader readout of virion interactions, extending beyond antibody binding to interactions with complement proteins, receptors, or other virions, provided that the interaction sufficiently alters particle properties (17, 18). Combined with quantitative bulk methods such as ddPCR, particle concentration measured by flow virometry can serve as a denominator for expressing bulk sample properties on a per-particle basis (**Fig. 4**). This approach can similarly be applied to measurements of viral protein abundance, incorporation of host proteins or other factors, lipid content, enzymatic activity, or infectivity. In addition, the rapid sampling of single-particle events characteristic of flow virometry enables real-time monitoring or endpoint quantification of dynamic changes, including those confined to rare subpopulations. We illustrated this capability by measuring lipid mixing between individual virus particles and liposome targets, both in real time and under antibody inhibition (**Fig. 5**). Broadly, any viral process that produces a measurable time-dependent change in fluorescence or light scatter could potentially be analyzed kinetically by the approach we describe here, including receptor engagement, antibody binding, particle aggregation or disaggregation, virion maturation, assembly or disassembly, and environmentally or pharmacologically induced transitions.

We further demonstrate that flow-based sorting can isolate virus subpopulations based on virion morphology, enriching filamentous particles from 20% to 65% of the population and retaining their infectivity for downstream analysis. Further enrichment may be achievable through successive rounds of sorting. We used a commercially available sorter and triggered events on V-SSC alone, eliminating potential inter-laser delay errors and associated inaccuracies in V-SSC-based size measurements (see Technical Note in the Supplementary Information). It is important to note that sorting results in a dilute product, so post-sorting concentration may be necessary depending on the application. The sensitivity of sorters is also generally lower than that of analyzers, making fluorescence-based labeling necessary for separating and sorting nanoparticles from noise; to avoid inhibition of infectivity by fluorescent antibody labeling, we engineered a MEDI8852-GFP construct designed to dissociate in the low pH environment of the late endosome (45). This antibody did not neutralize virus at the concentrations used for sorting (**Fig. 6B**) and, importantly, the sorting process did not alter per-particle infectivity (**Fig. 6E**). We thus established a workflow by which virion shape can be used to define and physically enrich virion subpopulations. This approach could be extended to other detectable particle features such as antigen abundance, binding state, or reporter fluorescence, allowing specific particle subpopulations to be isolated from heterogeneous samples. The sorted particles can then be used for downstream analysis of phenotypes such as infectivity, stability, immune recognition, or tropism.

Flow virometry enables single-particle analysis across many areas of virology, but several limitations should be considered. A single event measured by flow virometry is not necessarily an individual virion. An antigen-or a lipid-positive nanoscale event could correspond to an individual virion, an aggregate containing multiple virions, or another labeled particle such as an extracellular vesicle (EV); orthogonal validations of particle identity may be necessary. In our own hands, IAV particle counts and size distributions measured by flow virometry showed good agreement with orthogonal measurements, including HA-based assays and EM (2) (**Fig. 2C**). For PR8 IAV, we have found that HA-positive EVs are rare and form a distinct subpopulation when present at detectable levels. Flow virometry can also be limited by surface antigen abundance, labeling efficiency, and instrument sensitivity, which may restrict detection of dim or very small particles. The recent development of a designated small-particle flow analyzer, such as CytoFLEX Nano (Beckman-Coulter) might overcome many of these restrictions. However, we note that not all cytometers can be configured for real-time kinetic analysis; instruments which produce flow by pressurizing the sample chamber, for instance, take much longer to begin sampling and are limited to analysis of relatively slow phenomena or to endpoint measurements. Flow-based sorting has additional considerations. Given the lower sensitivity of current sorters, virus sorting generally requires fluorescent labeling. Because the labeling reagent itself can alter virion properties or infectivity, minimally perturbing labels are important when functional analysis of sorted populations is required. The non-neutralizing labeling reagent developed here illustrates one such approach. Downstream analysis of sorted populations is further complicated by particle dilution by cytometer sheath fluid, which may be improved by the use of narrower sorting nozzles. Despite these limitations, flow virometry and flow-based sorting provide complementary approaches for characterizing and isolating heterogeneous virus subpopulations. Broader adoption of nanoparticle flow cytometry in virology, together with the development of purpose-built nanoparticle flow cytometers, should accelerate research and further expand opportunities for studying virus population diversity and dynamics.

## Methods

### Cells, viruses and virus like particles

#### Cells

HEK293T, A549, MDCK-Siat1, and Vero cells were grown at 37°C and 5% CO_2_ to confluency before passaging. Cells were washed twice with HBSS then detached with 0.25% trypsin containing EDTA in HBSS. Growth medium was DMEM (HEK293T, A549, MDCK-Siat1) or EMEM (Vero), both supplemented with 10% FBS. L929 cells used in reovirus infections were grown in suspension culture at 37°C in spinner flasks. Cells were grown to 1-1.5 × 10^6^ cells/mL in Joklik’s MEM supplemented with 5% FBS and 2 mM glutamine and diluted to 2.5 × 10^5^ cells/mL at each passage. Expi293 cells were grown in Expi293 growth medium to 1-1.5 × 10^6^ cells/mL and diluted to 1-1.5 × 10^5^ cells/mL at each passage. Calu3 cells were grown to confluency in EMEM supplemented with 20% FBS at 37°C and 5% CO_2_ before passaging. Cells were washed and detached with accutase (Stemcell Technologies, Cat#07922).

HEK293T, A549, MDCK-Siat1, Calu3 cells were obtained from ATCC. L929 cells were gifted from Dr. John Parker’s group at Cornell University. Vero cells were gifted from Dr. Kartik Chandran at Albert Einstein College of Medicine.

#### Virus stocks and titration

IAV strains were propagated on MDCK-Siat1 cells for A/Puerto Rico/8/1934 (H1N1) and in Calu3 cells for all other strains, at an MOI of 0.002 in the presence of 1 µg/mL TPCK-treated trypsin (Sigma, Cat#T1426). Cells were monitored for cytopathic effect (CPE) development and virus was harvested approximately 48 hours post-infection. The supernatant was clarified by centrifugation at 1000 x g for 10 minutes. Viral infectivity was measured by TCID50 or image cytometry. For TCID50 assay, virus stocks were serially diluted four-fold in Opti-MEM containing 1 µg/mL TPCK-treated trypsin and used to infect MDCK-Siat1 cells in eight replicate wells per dilution. At 4–7 days postinfection, wells were evaluated for CPE, and TCID50 titers were calculated using the Reed–Muench method (46). To measure viral infectivity by image cytometry, samples were serially diluted and used in infections of A549 cells. To prevent spread, TPCK-treated trypsin was omitted and 20 mM NH_4_Cl added 4 hours post infection. At 20 hpi, samples were washed with PBS and fixed with 4% PFA for 20 min, then washed again with PBS. HA and nuclei were then stained for 1 h at room temperature in PBS containing 1% BSA using 50 nM MEDI-scFv-sfGFP antibody (20) and 4 µM Hoechst 33342 (Thermofisher, Cat#62249), respectively. Samples were then imaged on a Cytation 5 (Agilent) and analyzed using CellProfiler (47) to count HA-positive and HA-negative cells. The fraction of infected cells was calculated by dividing the number of HA-positive cells by the total cell count per well and related to particle input to determine the number of particles per infectious unit.

VSV Indiana serotype seed stock was a gift from Dr. Jonathan Yewdell’s group at the National Institutes of Health and was propagated according to the published protocol (48). Briefly, a flask of 90% confluent Vero cells was infected at an MOI of 0.0001 for 1 hour in EMEM without FBS. The medium was then replaced with EMEM with 10% FBS. The flask was incubated at 37 °C in 5% CO_2_ for 48 hours. The virus was harvested and centrifuged at 2,900 × g at 4 °C for 10 min, and the supernatant was collected. To generate the reovirus stock, spinner-adapted L929 cells (2 × 10⁸ cells) were incubated with type 1 Lang reovirus at an MOI of 5 in 10 mL Joklik’s MEM (JMEM) containing 5% FBS for 1 hour at room temperature. After adsorption, the cells were resuspended in JMEM with FBS to a final volume of 400 mL and maintained in spinner culture at 35 °C until the viable cell count was reduced to 65%, as assessed by trypan-blue staining. Cells were harvested by centrifugation, resuspended in 8 mL HO buffer (250 mM NaCl, 10 mM Tris, pH 7.4, and 10 mM β-mercaptoethanol), and frozen. After thawing, deoxycholate was added to a final concentration of 0.1%, and the suspension was incubated on ice for 30 minutes. Vertrel XF (5 mL) was added, and the sample was sonicated to disrupt the cells. The resulting emulsion was centrifuged at 10,000 × g for 10 minutes. The aqueous phase was collected, sonicated with a further 5 mL Vertrel XF, and centrifuged again at 10,000 × g for 10 minutes. The aqueous phase was layered onto a 1.25-1.45 g/mL CsCl gradient and centrifuged at 100,000 × g for 2 hours. The primary band was collected and extensively dialyzed against HVB (150 mM NaCl, 10 mM HEPES, pH 7.4, and 10 mM MgCl_2_) to generate the viral stock.

EBOV VLPs were produced by transfection using the Expi293 Expression System (Thermofisher, Cat#A14635) according to the manufacturer’s protocol. VLPs were generated using the following plasmid ratios: 0.48 VP40-GFP:0.053 WT-VP40:0.13 full-length GP: 0.33 NP. Two days post transfection, supernatant was clarified by centrifuging for 10 min at 1000 × g.

CoV NL63 was obtained from BEI Resources (NR-470). It was propagated in Vero cells at an MOI of 0.003 PFU/cell, then purified sequentially through a 20% sucrose cushion and a 20–60% continuous sucrose gradient in HNE20. Viral titers were determined by plaque assay on Vero cells. VACV Western Reserve strain (tk-) was gifted from Dr. Kartik Chandran at Albert Einstein College of Medicine. RSV strain A2 was gifted from Dr. Nicholas Guydosh’s group at National Institutes of Health. A/Texas/37/2024 (H5N1) was gifted from Dr. Jeffery Taubenberger’s group at the National Institutes of Health. Experiments involving H5N1 were performed under biosafety level 3 (BSL-3) conditions at the C.W. Bill Young Center BSL-3 in Bethesda, Maryland, USA. All work with H5N1 was approved under NIH pathogen registration PRD-24-BMC-179. All experiments with biosafety level 2 (BSL-2) IAV strains, VSV, EBOV-VLPs, and RSV were approved under NIH pathogen registration RD-23-IV-10, with VACV under NIH pathogen registration 8856, and with reovirus under registration RD-23-IV-11. NL63 experiments were performed at Brandeis University under the pathogen registration 16004.

### Fluorescent labeling reagents

#### MEDI8852 derived antibody design

To generate a small fluorescent label that could access the conserved base of IAV HA in virometry, we generated a single-chain variable fragment with the moxGFP inserted between the heavy and light chains, which was synthesized by Biointron. The overall construct organization was Signal peptide – VL – SGG linker – moxGFP – GGSSSG linker – VH – 6xHis. See supplement information for the antibody sequence.

The design of other MEDI ScFv constructs, including WT MEDI-ScFv-eGFP and MEDI-scFv-sfGFP used for infectivity assays, has been previously reported (20). To reduce the neutralization activity of WT MEDI-scFv-eGFP, residue 31 of MEDI-ScFv-eGFP was selected for histidine mutagenesis using the SIpHAB tool (45) and mutated to histidine using Gibson-assembly (FWD primer: CCGTCTCCATCTACAACGCCGT, reverse primer: GTTGTAGATGGAGACGGAGTCGC), yielding H31 MEDI-scFv-eGFP.

#### Antibody expression and purification

Expi293 cells were transfected with plasmids expressing the specified MEDI scFv construct using expifectamine transfection reagent (ThermoFisher, Cat#A14524) according to manufacturer’s protocol. Five days post transfection, culture supernatants were clarified of cell debris by centrifugation and filtering through a 0.45 µm filter. Supernatants were combined with imidazole to a final concentration of 2 mM for MEDI8852 scFv-moxGFP or 0.2 mM for MEDI scFv-eGFP and purified over a 5 mL HisTrap column (Cytiva, Cat#17524801) for MEDI scFv-moxGFP or Nickel NTA-Agarose column (Qiagen, Cat#30210) for MEDI scFv-eGFP2. Each antibody sample was further purified by size exclusion chromatography on a Superdex 200 increase 10/300 column (Cytiva, Cat#28990944) equilibrated in PBS to remove protein aggregates that might contribute to noise and/or promote virion aggregation in flow virometry. MEDI-scFv-sfGFP was purified as previously described (20).

#### Antibody fluorescent conjugation

Lyophilized NHS ester dyes (ThermoFisher, Cat#s 62266, 62263, 46403, 46401) were resuspended in anhydrous dimethyl sulfoxide (DMSO). Dye solution was combined with antibodies at a molar ratio ranging from 20:1 to 100:1 in PBS + 0.1 M sodium bicarbonate. Dye:target ratio may need to be adjusted on a case-by-case basis to optimize fluorescent signal. Volume of DMSO in the reaction mixture should not exceed 10%. Reactions were performed for 1 hr at room temperature in the dark. Unbound dye was removed from labelled antibodies using Harvard Apparatus Macro SpinColumns with G25 or G50 resin according to manufacturer instructions (Harvard Apparatus, Cat#s 74-3901 and 74-3902).

### Sample preparation and flow virometry acquisition

#### Optical alignment for nanoparticle detection

To produce the scattered-light and fluorescence signals required to resolve single virus particles, engineers from the instrument manufacturer optimized laser alignment for nanoparticle detection. A sample of fluorescently labelled virions was continuously analyzed as the positioning of the focusing lens was tuned for each laser. Real-time correlation between changes in alignment and changes in the amount of detected light corresponding to nanoparticles was key to optimizing detection for our instrument (Cytoflex S), and we expect that a similar strategy will be useful for other cytometry platforms as well. We reiterate here that the 405-nm laser substantially outperforms the 488-nm laser for virus particle detection by side-scatter; however, the 488-nm laser may be sufficient for detection of larger nanoparticles if proper optical alignment is achieved. We recommend use of Beckman-Coulter nanoViS low mix beads (Beckman Coulter, Cat# D03231) as a calibration standard to verify small particle capability of a cytometer. We found that the 80-nm diameter polystyrene bead standard produced a similar V-SSC signal to that of IAV particles (**Fig. S1A**).

#### Flow virometry with antibody or lipid staining

For antibody labeling, 2 µL of the virus sample was mixed with 2 µL of antibody at a final concentration of 20 nM for antibodies targeting accessible epitopes (e.g., an HA head antibody), or 200 nM for antibodies targeting less accessible epitopes (e.g., an HA stem antibody) and incubated for 1 h at room temperature. The 96-well cytometry plates (BRANDTECH, Cat# 781601) were treated with 0.1% BSA in PBS for at least 5 min to prevent nonspecific binding of virions to the plastic (**Fig. S2**). Plates were then rinsed with water to remove excess BSA and dried before adding sample. 1.2 µL of virus/antibody mixture was added to 300 µL of HNE20 (20 mM HEPES, 150 mM NaCl, 1 mM EDTA, pH 7.4) in the 96-well plate. All samples were run either on CytoFLEX S or CytoFLEX LX flow cytometers (Beckman Coulter). Before acquiring sample data, clean buffer was run for approximately 5 minutes to clear out sample lines. Sample acquisition was performed at a flow rate of 10 µL/min. Ensure that the flow rate of the machine is regular and consistent before measurement both by using built in cytometer flow rate calibration protocols and by monitoring acquired events/s to ensure stable flow (**Fig. S1B**).

When performing flow virometry, sample carryover may occur, in which a measurement is contaminated by virions from the previous sample (**Fig. S1C**). A 20-second backflush between samples can help reduce carryover. Carryover is particularly problematic when a low-titer sample is measured directly after a high-titer sample. Low-titer samples should be measured before high-titer samples when possible. Otherwise, running clean buffer or water after a sample can help in clearing instrument lines of residual virions. For persistent carryover, it may even be necessary to run a cleaning agent, followed by water, to fully purge the instrument of leftover sample. For these purges, we ran Beckman-Coulter’s FlowClean cleaning agent (Beckman Coulter, Cat#C48093) for 60s followed by Milli-Q water for 300s.

Each sample was mixed for 5 seconds before data acquisition. Event acquisition was triggered on V-SSC height. The trigger threshold was empirically determined by starting at a high value and progressively lowering it until the entire fluorescent virion population was captured. The highest threshold that captured the full virion population was used to minimize acquisition of background noise (**Fig. S1D**). Because cytometer noise is elevated at the start of sample acquisition, events acquired during the first 120 s are typically excluded from analysis **(Fig. S1B).** With sufficient labeling, virion events are fully separated from noise in the fluorescent channel, allowing the virion population to be gated accordingly.

For lipophilic dye labeling, a 1 mM R18 (Thermo Fisher, Cat# O246) stock solution in ethanol was diluted in HN20 buffer to a final concentration of 10 µM. It is important to ensure that the final diluted dye contains no more than 1% ethanol by volume. Purified virus aliquots were made at approximately 1.2 mg/mL total protein concentration as assessed by the Bradford assay. 20 µL of 10 µM R18 was then mixed with 5 μL of purified IAV (6 μg of total viral protein). Mixtures were incubated at room temperature in the dark for 60 mins. After incubation, a PD-10 desalting column (Cytiva, Cat#17085101) was used to separate virions from free dye. Virion concentration post labeling and labeling efficiency were then measured by flow virometry.

### Particle counting and morphology analysis

Purified PR8 IAV was diluted 100-times before being subjected to seven additional 10-fold serial dilutions. The unlabeled sample was diluted 1:20 in HNE20 buffer before measurement of particle titer, with 2000 A.U. in VSSC height used as the gate boundary separating virion events from background noise. 2 × 10^5^ total events were recorded in the unlabeled virion gate. Labeled sample was bound 1:1 with 400 nM MEDI-scGFP before addition of 1.2 µL labeled sample to 300 µL HNE20 for acquisition. 2.5 × 10^4^ events were acquired in the labeled virion gate.

### Antibody binding and C1q induced aggregation

#### Size shift of virus particles after antibody binding

Purified PR8 IAV was diluted to approximately 10^6^ particles/µL and mixed 1:1 with 200 nM antibody. After 1 hr at room temperature, samples were diluted 1:20 in HNE20 and subjected to unlabeled flow virometry. Head-binding antibodies were H36-26 (Sb antigenic site) (24), Y8-1A6 (Sa antigenic site) (49), H28-D14 (Sb antigenic site) (24), and H35-C12 (Cb antigenic site) (50). Stem-binding antibodies were CR9114 (51), MEDI8852 (29), Fi6 (52), and 16G8 (53).

#### C1q-antibody aggregation of virions

Purified PR8 IAV was diluted to approximately 10^6^ particles/µL and combined 1:1 with 200 nM antibody H28-D14, or with a mixture of 200 nM antibody H28-D14 and 140 µg/mL human C1q. After 1 hr at room temperature, samples were diluted 1:20 in HNE20 and subjected to unlabeled flow virometry.

### Genome to particle and defective genome measurements

#### Inducing aggregation by NA inhibitor

MDCK-Siat1 cells were infected at MOI 0.6 IU/cell. At four hours post-infection (hpi), the media was supplemented with 1:1000 dilution of bacterial neuraminidase (from Vibrio cholerae, Sigma-Aldrich, Cat#11080725001) 4 hpi. A final concentration of 1000 nM of oseltamivir acid (LGC Standards *née* Toronto Research Chemicals, Cat#O700980) was added at 22 hpi. The supernatant was collected at 24 hpi for analysis.

#### Measuring genome-to-particle ratio by flow virometry and ddPCR

To quantify genomes in an IAV sample, 10 µL of a known concentration of VSV was mixed with 130 µL of the sample. A final concentration of 50 ng/mL of RNaseA was added to the virus mixture and incubated at 37 °C for 30 min to degrade any free RNA present in the sample. Viral RNA extraction was performed following the manufacturer’s instructions (QIAamp Viral RNA Kits, Qiagen, Cat#52906). Reverse transcription reaction used M-MuLV reverse transcriptase (M-MuLV Reverse Transcriptase, New England Biolabs, Cat# M0253L) according to the manufacturer’s protocol with primers: Uni-12 (5’-ACGCGTGATCAGCAAAAGCAGG-3’), VSV-Tr-F (5’-AAATCATGAGGAGACTCCAAAC-3’). The viral genome cDNA was then 10-fold serially diluted 50 to 50000 times and used as the template for ddPCR: 11 µL 2X EvaGreen Supermix (QX200 ddPCR EvaGreen Supermix, Bio-rad, Cat#186-4035), 0.7 µL each forward and reverse primers (10 µM, Table 1), 5.5 µL cDNA, 4.1 µL nuclease-free water. The reaction was then performed with the QX200 Droplet Digital PCR system (Bio-rad) following the manufacturer’s instructions. The IAV genome-to-particle ratio was calculated based on the raw genome copy concentration obtained from ddPCR multiplied by the cumulative dilution factor and divided by the VSV genome-to-particle ratio to adjust gain/loss over the process.

**Table 1.**
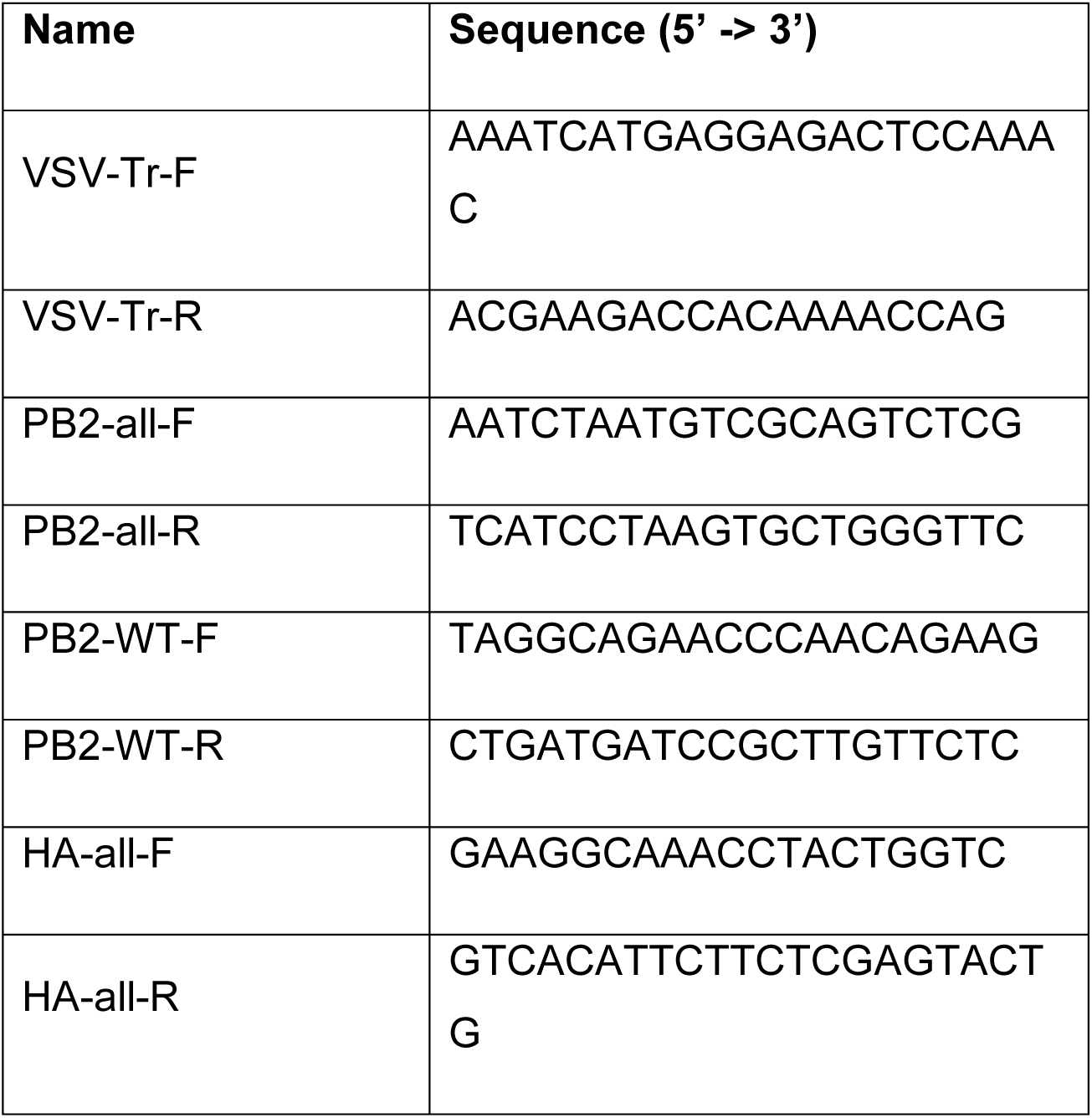
Primers used for ddPCR.

### Single-pair Cytometric Analysis of Lipid Mixing

#### Virion labeling and preparation

R18-labeling concentration is selected to provide sufficient fluorescence for virion detection by flow virometry before lipid mixing and sufficient dequenching to distinguish lipid-mixed from unmixed virions. The extent of fluorescence increase also depends on the amount of available membrane area in the target; small targets do not permit the same extent of R18 dilution as the larger ones. We labeled purified, sphere-enriched XUd virions with 4 μM R18; the final ethanol concentration in the labeling mixture was 1 % by volume. R18-labeled virions were frozen in single-use aliquots at-80 °C; they were thawed rapidly and used within one day of thawing. Their concentration after thawing was determined by flow virometry.

#### Liposome preparation

We prepared liposome targets as described in (Planitzer et al., 2026); briefly, chloroform-dissolved DOPC (1,2-dioleoyl-sn-glycero-3-phosphocholine, Avanti Research, Cat#850375), POPC (1-palmitoyl-2-oleoyl-glycero3-phosphocholine, Avanti Research, Cat#850457), cholesterol (Avanti Research, Cat#700100), fluorescein-DHPE (N(Fluorescein-5-Thiocarbamoyl)-1,2-Dihexadecanoyl-sn-Glycero-3-Phosphoethanolamine, Triethylammonium Salt, Invitrogen, Cat#F362), and biotin-PE (1,2-dioleoyl-sn-glycero-3phosphoethanolamine-N-(cap biotinyl) (sodium salt), Avanti Research, Cat#870273) and ethanol-dissolved DiD (1,1’-Dioctadecyl-3,3,3’,3’-Tetramethylindodicarbocyanine, 4-Chlorobenzenesulfonate Salt, Invitrogen, Cat#D7757) were mixed at a 40:40:20:0.025:0.0025:0.0452 molar ratio. The mixture was dried first using a stream of pure argon gas and then further by vacuum dessication for at least two hours. The lipid film was suspended at 9.24 mg/mL in 20 mM HEPES pH 7.4, 150 mM NaCl, 10 μM CaCl2 containing lipid-ssDNA conjugate (sequence: /5Lipid/5’-TCG ACA CGG AAA TGT TGA TGT TGA-3’) at 1 % total lipid. The lipid-ssDNA conjugate was prepared in-house as described (21) and presents a platform on which complementary synthetic ssDNA-receptor conjugates may be hybridized. An alternative option for IAV fusion studies is the widely employed ganglioside GD1a which presents α2,3-linked sialic acids, is readily commercially available, and may be incorporated in the lipid mixture before drying into a lipid film (54). The crude liposome suspension was shaken at 37 °C for 30 minutes, subjected to five freeze-thaw cycles between liquid nitrogen and warm water with vortexing between cycles, and extruded through a 1-μm filter 30 times. Liposomes were degassed under house vacuum for 10 minutes, stored at 4 °C in a parafilmed microcentrifuge tube filled with argon gas, and used within two months. For spCALM experiments, the liposomes were counted by flow virometry using DiD fluorescence and violet side-scatter intensity to resolve from background noise.

#### Performing spCALM

Before spCALM, R18-labeled XUd virions were incubated with MEDI-Fab for one hour at room temperature at the indicated concentrations. Separately, liposomes were incubated with a 6:4 mixture of a custom-synthesized 3’sialyllactose-ssDNA conjugate (sequence: /6’SL/5’-TCA ACA TCA ACA TTT CCG TGT CGA-3’) and non-receptor-ssDNA (sequence: 5’-TCA ACA TCA ACA TTT CCG TGT CGA-3’) and with MEDI-Fab for one hour at room temperature. 1.67e6 virions were combined with 2e6 liposomes in 20 μL and shaken at 34 °C for five minutes. 80 μL of pre-warmed HN20C (20 mM HEPES pH 7.40, 140 mM NaCl, 10 μM CaCl_2_) was then added, and sample acquisition by the flow cytometer was initiated at a flow rate of 80 μL/min. After 10 seconds, 900 μL of pH 5.68 buffer (30 mM MES, 30 mM acetic acid, 140 mM NaCl, 10 μM CaCl2) was added to the sample tube and mixed; samples were acquired for a total of 10 minutes each. The cytometer tube holder was filled with water and wrapped with a heating strip held at 34 °C during the lipid mixing reaction. To improve temporal resolution, virion transit time through the cytometer fluidics was minimized by using a higher flow rate than for the other flow virometry measurements described here. Virion and liposome concentrations were adjusted to avoid excessive abort rates at the higher flow rate.

#### Statistical analysis

Single virion-target pair data were gated and exported using FlowJo (v10.9.0; Becton, Dickinson & Company); lipid-mixing kinetics of these pairs were then quantified using custom MATLAB (R2023a; The Mathworks) software, as described in (21). Briefly, characteristic reference R18 fluorescence distributions of unfused and lipid-mixed pairs were estimated by fitting Burr distributions within the pre-pH-drop time interval (for unfused) and late time reaction plateau (lipid-mixed); the late time plateau represented a mixed population of unfused and lipid-mixed pairs, so the characteristic lipid-mixed distribution was estimated by gaussian mixture model deconvolution. For each sample, detected pairs were binned so that each bin contained approximately the same number of events; an expectation-maximization algorithm was used to estimate the abundance of the lipid-mixed distribution relative to the unfused distribution within each time bin to produce a kinetic trajectory of lipid mixing. Lipid mixing efficiency is estimated by the difference in lipid-mixed fraction between the late-time plateau and pre-pH-drop interval. pH drop time is detected from the maximum rate of change of detected particle concentration from low-pH buffer addition; the lipid-mixing lag time is the time at which an isotonic fit of the lipid-mixed fraction trajectory reaches half of the maximal value, less the pH-drop time. Linear and 4-parameter logistic fitting were performed using Graphpad Prism (v10.6.1, GraphPad Software).

### Electron microscopy, virion sorting and infectivity

#### Comparison of virus size as measured by EM and flow virometry

12 hpi, virus-containing supernatant was harvested from cells and replaced with OptiMEM without TPCK trypsin. 20 hpi, virus-containing supernatant was harvested and both supernatants were clarified by centrifugation at 1000 x g for 10 minutes. Samples were stained for flow virometry by mixing 1:1 with 25 nM DyLight 550-labeled (Thermofisher, Cat#62262) HC19 antibody (55), incubating for 1 hour at room temperature, then diluting in HNE20. Samples were analyzed using a Cytoflex S flow analyzer (Beckman Coulter) as described above.

For EM, samples were purified over a 20-60% sucrose step cushion prepared in PBS at 100,000 x g for 1 hour in a SW28 rotor (Beckman Coulter). Virus samples were collected at the 20-60% sucrose interface and fixed at 1:2 ratio of sample to 4% paraformaldehyde for 30 minutes at room temperature. Samples were then diluted in PBS and further concentrated over a 30-60% sucrose step cushion by spinning at 100,000 x g for 1 hour in a SW28.1 rotor (Beckman Coulter). To further concentrate virus to necessary levels for electron microscopy, samples were concentrated on a 60% sucrose cushion in a TLS55 rotor (Beckman Coulter) at approximately 50,000 x g for 1.5 hours. Samples were then diluted 1:2 with PBS and further concentrated over a 60% sucrose cushion in a TSL55 rotor at approximately 50,000 x g for 1.5 hours to a final volume of 20 µL. Samples were applied onto EM grids, stained with 2% phosphotungstic acid as previously described (56). Micrographs were acquired on a Talos L120C (Thermo Fisher), operating at 120 kV, with a Rio camera (Gatan) using SerialEM (57). Particle sizes were measured in FIJI (58).

#### Antibody neutralization assay for WT and H31 MEDI-scFv-eGP2 constructs

XUd was labeled with the indicated antibodies by incubating clarified viral supernatant 1:1 with 2X the indicated concentration for 1 hour at room temperature (RT) or with antibody dilution buffer alone (HNE20 + 0.02% BSA). Virus particles were then used to infect A549 cells at an MOI of 0.1, and infectivity was assessed via image cytometry as described above.

#### Labeling and sorting of IAV

XUd was labeled by incubating clarified viral supernatant 1:1 with 500 nM of H31 MEDI-scFv-eGFP for a final labeling concentration of 250 nM for 1 hour at RT. Labelled virus was further diluted 1:250 in HNE20 buffer. Samples were sorted on a Cytoflex SRT (Beckman Coulter, Brea CA) using 1X PBS as sheath fluid and a 100 µm nozzle. Virus events were detected using a VSSC-H trigger of 300 and gated first on VSSC-A versus FITC-area and then on VSSC-H and FITC-height (**Fig S4**). Size-specific gates used for sorting were established on VSSC-A.

#### Infectivity of sorted virus

A549 cells were spinoculated with 1 mL of each sorted fraction for 1 hour at room temperature at 2300 x g. Following spinoculation, inoculum was removed from cells and replaced with OptiMEM without TPCK trypsin. The cells were then incubated at 34 °C for 20 hours and infectivity was calculated by image cytometry as described above.

## Acknowledgements

We thank John Parker and his group at Cornell University for providing L929 cells; Kartik Chandran and his group at Albert Einstein College of Medicine for providing Vero cells and vaccinia virus samples; Jonathan Yewdell and his group at the National Institutes of Health (NIH) for providing the vesicular stomatitis virus seed stock; Max Nibert and his group at Harvard University for providing the reovirus seed stock; Nicholas Guydosh and his group at NIH for providing respiratory syncytial virus; Robert Davey and his group at Boston University for providing EBOV NP plasmid, Kartik Chandran’s group at Albert Einstein College for providing EBOV VP40, EBOV VP40-GFP, and EBOV GP plasmids, and Jeffery Taubenberger and his group at NIH for providing IAV Texas/37/2024 (H5N1). We thank BEI Resources for providing human coronavirus NL63 (NR-470).

We thank Iyadh Douagi, Melanie Cohen and Julie Laux at the Cytometry Section, Research Technology Branch, NIAID, NIH for assistance with sample acquisition under BSL3 conditions at the C.W. Bill Young Center in Bethesda, Maryland. We thank Kyra Kerkhofs at NIH for her collaboration in developing flow virometry for RSV and conducting experiments with Antonio Santos Del Peral. We thank Jia Niu at the Department of Chemistry, Boston College for providing the oligonucleotide-conjugated receptors and lipids used in this study. We also thank Vinod Nair and Cindi Schwartz at the Electron Microscopy Unit, Research Technologies Branch, NIAID, NIH for assistance with electron microscopy.

This research was supported in part by the Intramural Research Program of the National Institutes of Health (NIH). The contributions of the NIH author(s) are considered Works of the United States Government. The findings and conclusions presented in this paper are those of the author(s) and do not necessarily reflect the views of the NIH or the U.S. Department of Health and Human Services.

## Declaration of Interests

T.I. is involved on a patent related to the flow virometry methodology: PCT/US2022/042125, status pending. The authors declare no other competing interests.

## Supplementary Figures

## Supplementary Information

### Technical note: Measurement delay due to fluidic instability in the Cytoflex platform

Many, but not all, flow cytometers and sorters, including the Cytoflex S and SRT used here, use an optical configuration in which individual particles pass through different lasers sequentially. To associate measurements acquired from the same particle as it passes through different lasers at different times, the cytometer establishes the inter-laser delays during daily quality control and adjusts them based on the selected flow rate. To acquire data, a triggering threshold is set on one or more of the detection channels associated with one or more lasers. When a particle generates signals that satisfy the specified triggering criteria, an event is logged. The cytometer then acquires signals from the different detection channels at times that account for the previously determined inter-laser delays.

When the flow rate is stable, this scheme enables signals from individual particles to be accurately measured across all four lasers. However, when the flow rate changes unexpectedly (e.g., because of bubble formation in the fluidic path), the calculated delays become misaligned with particle transit through the non-triggering lasers, causing signal acquisition to occur before or after the peak and instead sample its shoulder. This results in reduced measured intensity in channels associated with the non-triggering lasers and can render these measurements unreliable unless flow stability is carefully monitored and the fluidic path is kept free of air bubbles. Importantly, these delay issues do not affect the channel used in triggering because signal acquisition is synchronized to the particle crossing the trigger threshold, ensuring that the full signal is captured. For this reason, when analysis relies on quantitative signal intensity in a particular detection channel, as when distinguishing IAV filaments from spherical virions by V-SSC, it is best to trigger on a channel associated with the same laser.

To maintain reliable V-SSC measurements for distinguishing small and large virions, we therefore use V-SSC to trigger event detection. With sufficiently bright fluorescent antibody labeling, virions remain distinguishable from background noise even when a suboptimal inter-laser delay reduces the measured fluorescence intensity, allowing reliable gating of the virion subpopulation. The virion-gated events are then classified as small or large using a fixed V-SSC threshold applied across samples.

We further note that we have not experienced this delay issue during measurement of larger objects, such as cells. We hypothesize that flow instability produces a relatively small spatial offset between the optimal measurement position and the actual particle position. For cells, this offset may be small relative to the dimensions of the object, allowing the signal to be captured despite the timing error.

### scMEDI-GFP design

Signal peptide – VL – SGG linker – moxGFP – GGSSSG linker – VH – 6xHis:

**MHSSALLCCLVLLTGVRA-**

**DIQMTQSPSSLSASVGDRVTITCRTSQSLSSYTHWYQQKPGKAPKLLIY AASSRGSGVPSRFSGSGSGTDFTLTISSLQPEDFATYYCQQSRTFGQG TKVEIK-SGG-**

**SVSKGEELFTGVVPILVELDGDVNGHKFSVRGEGEGDATNGKLTLKFIC TTGKLPVPWPTLVTTLTYGVQCFSRYPDHMKRHDFFKSAMPEGYVQE RTISFKDDGTYKTRAEVKFEGDTLVNRIELKGIDFKEDGNILGHKLEYNF NSHNVYITADKQKNGIKANFKIRHNVEDGSVQLADHYQQNTPIGDGPVL LPDNHYLSTQSKLSKDPNEKRDHMVLLEFVTAAGITHGMDELYK-GGSSSG-**

**TQVQLQQSGPGLVKPSQTLSLTCAISGDSVSSYNAVWNWIRQSPSRGL EWLGRTYYRSGWYNDYAESVKSRITINPDTSKNQFSLQLNSVTPEDTA VYYCARSGHITVFGVNVDAFDMWGQGTMVTVSS-HHHHHH**

---

**Figure S1:**
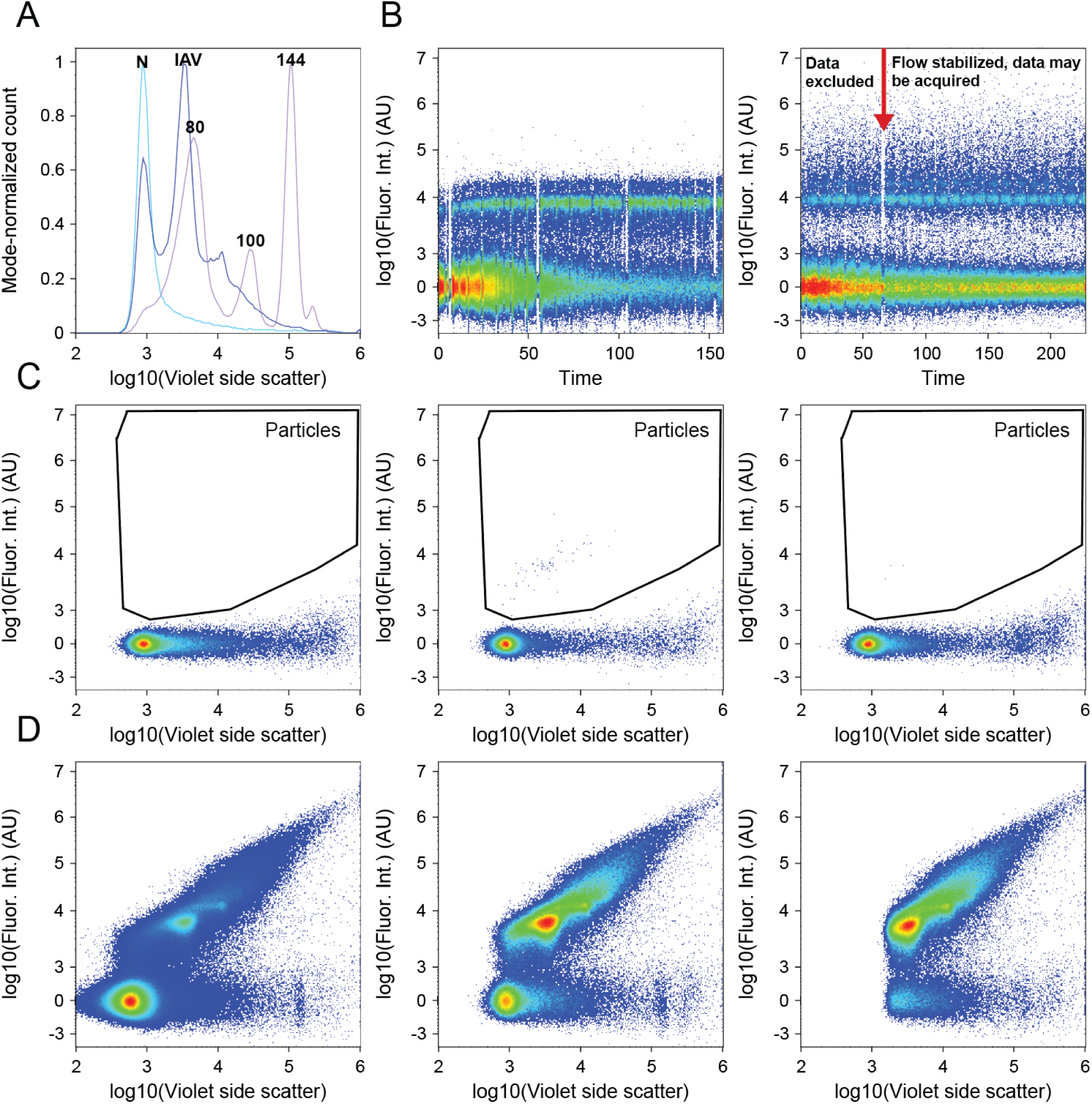
Technical considerations in flow virometry. (A) Overlay of the V-SSC signal from the buffer alone (*light blue*), Beckman-Coulter nanoVIS beads at 80 nm, 100 nm, and 144 nm (*magenta*), and IAV virions (*purple*). Peaks are labeled for noise (N), bead diameters (*numerical, nm*) or virions (IAV). (B) Comparison of event acquisitions with unstable (*left*) or stable (*right*) flow. Unstable flow necessitates a change of machine tubing or other repairs. (C) HNE20 buffer at the beginning of a run (*left*), HNE20 buffer acquired directly after a high-titer sample of IAV (*center*), and HNE20 buffer after a FlowClean/water rinse (*right*). Carryover of sample signal is reduced after the extra rinse. For most cases, water or buffer alone can reduce carryover. We recommend using FlowClean or other detergents sparingly to avoid introducing bubbles, which increase noise. (D) IAV sample acquired with varying V-SSC triggering thresholds. A low threshold (*left*) captures sub-virion fluorescent events, but a high level of noise events increases the abort rate. A ‘middle’ threshold (*center*) loses a small amount of sub-virion signal but has much lower noise. A high threshold (*right*) has low noise but cuts off a significant portion of virion events.

**Figure S2.**
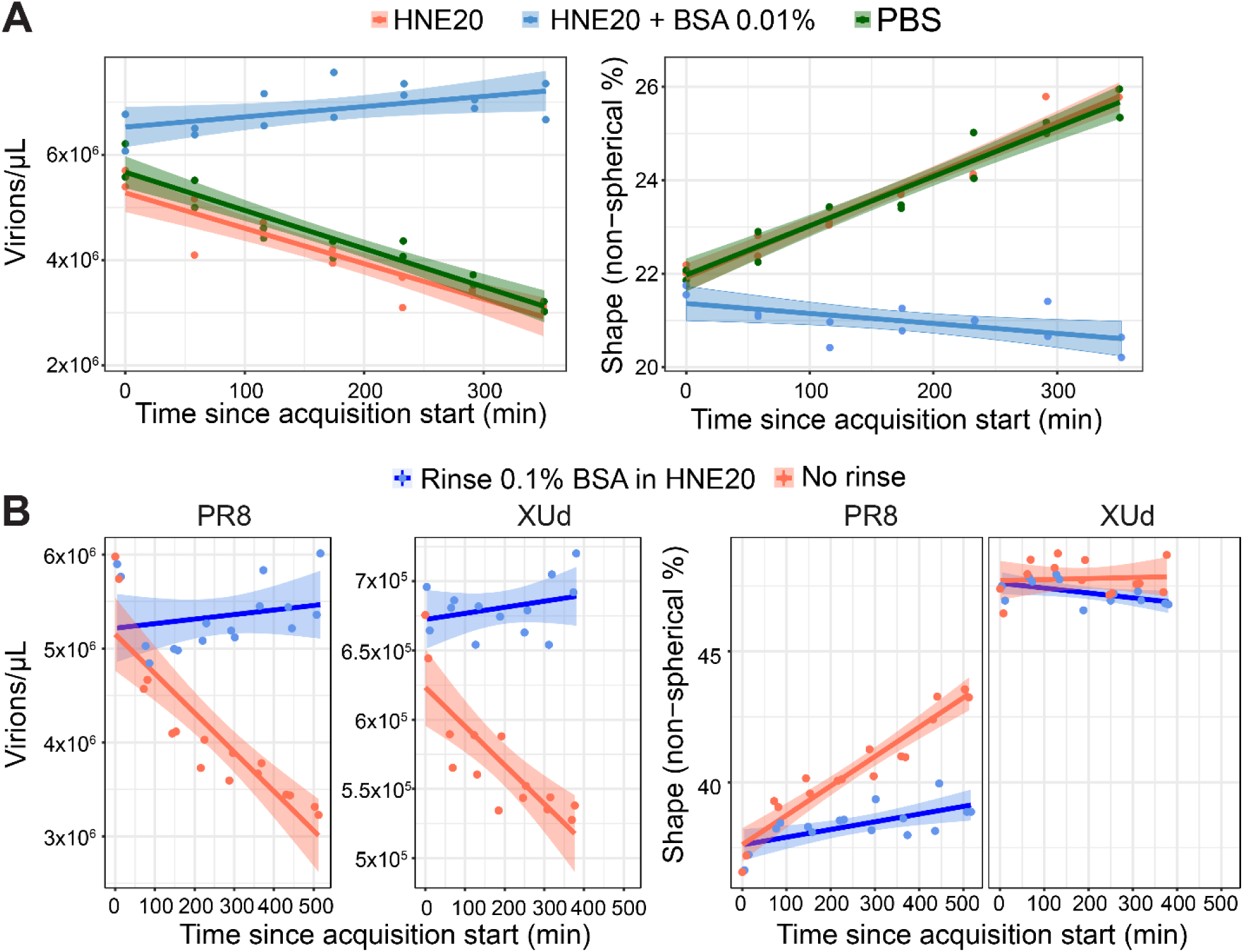
BSA reduces sample loss during flow virometry acquisition. (A) PR8 samples were acquired by flow virometry, and particle concentration was plotted over acquisition time. Samples were diluted in HNE20, HNE20 supplemented with 0.01% BSA, or PBS to assess buffer-dependent changes in particle recovery and shape. (B) PR8 and XUd samples diluted in HNE20 were acquired from plates that were either untreated or pretreated with 0.1% BSA and then rinsed twice with HNE20.

**Figure S3.**
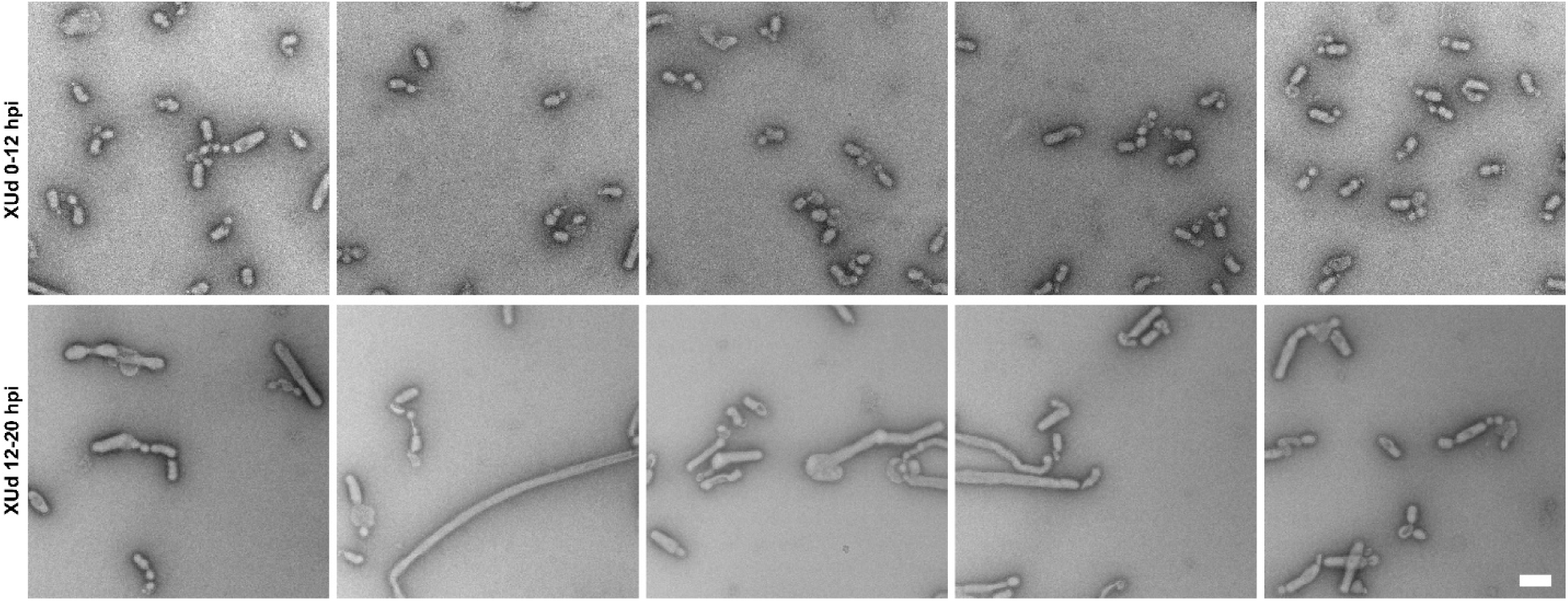
XUd produces increasingly filamentous virions at later stages of infection. *Temporal changes in XUd virion lengths revealed by electron microscopy (EM)*. Representative negative-stain electron micrographs of XUd virions produced by 12 h post-infection (*top*) and 12–20 h post-infection (*bottom*) from infection of MDCK-Siat1 cells. Scale bar = 200nm.

**Figure S4:**
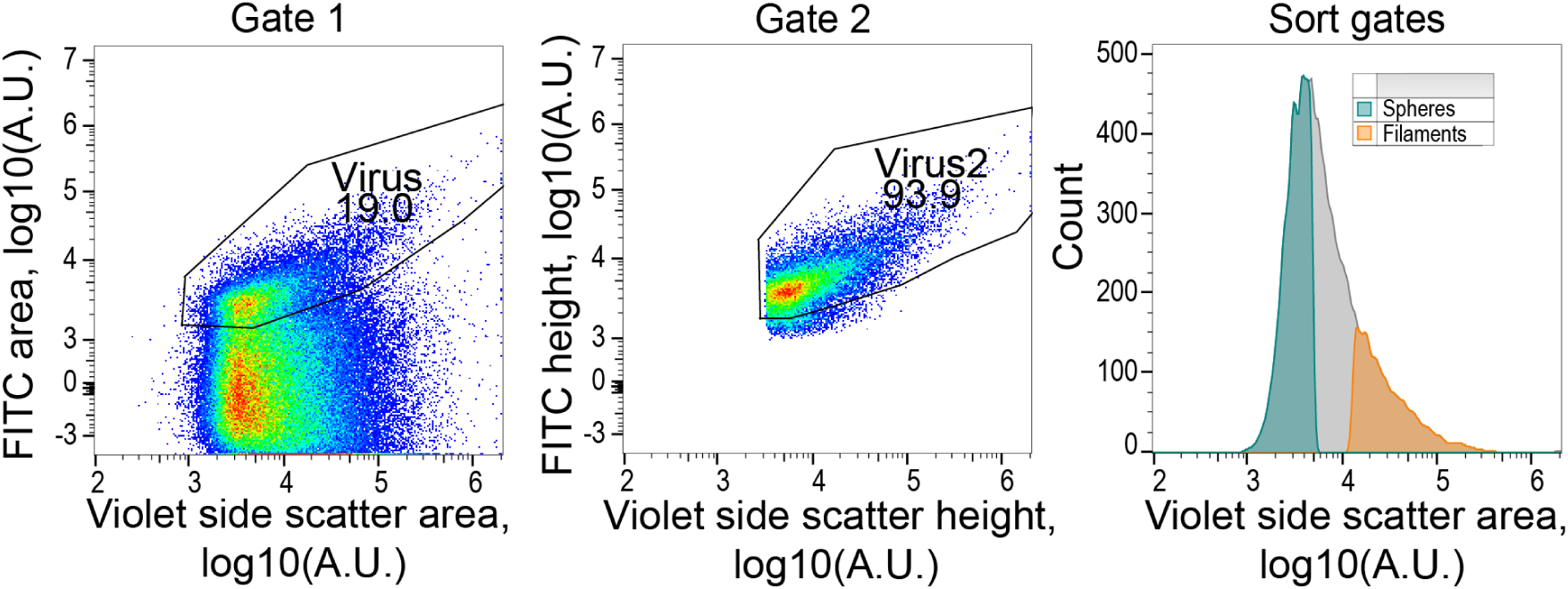
Gating strategy to separate virions from noise on the sorter: XUd stained with 250 nM of H31 MEDI scFv-eGFP, gated first on FITC-Area and VSSC-Area (*left*) then on FITC-Height and VSSC-Height (*middle*) to better isolate the virion population from inherent noise on the sorter prior to establishing sort gates (*right*) in VSSC-Area.

